# Single-molecule long-read sequencing reveals a conserved selection mechanism determining intact long RNA and miRNA profiles in sperm

**DOI:** 10.1101/2020.05.28.122382

**Authors:** Yu H. Sun, Anqi Wang, Chi Song, Rajesh K. Srivastava, Kin Fai Au, Xin Zhiguo Li

## Abstract

Sperm contributes diverse RNAs to the zygote. While sperm small RNAs have been shown to be shaped by paternal environments and impact offspring phenotypes, we know little about long RNAs in sperm, including mRNAs and long non-coding RNAs. Here, by integrating PacBio single-molecule long reads with Illumina short reads, we found 2,778 sperm intact long transcript (SpILT) species in mouse. The SpILTs profile is evolutionarily conserved between rodents and primates. mRNAs encoding ribosomal proteins are enriched in SpILTs, and in mice they are sensitive to early trauma. Mouse and human SpILT profiles are determined by a post-transcriptional selection process during spermiogenesis, and are co-retained in sperm with base pair-complementary miRNAs. In sum, we have developed a bioinformatics pipeline to define intact transcripts, added SplLTs into the “sperm RNA code” for use in future research and potential diagnosis, and uncovered selection mechanism(s) controlling sperm RNA profiles.

## Introduction

Sperm contribute DNA and diverse species of RNA to the zygote during fertilization^1–4^. While sperm RNA contribution is relatively small compared to oocyte RNA, they mediate transgenerational inheritance^5–11^. While this form of epigenetic inheritance is well established in plants, fungi, worms, and flies^12^, we are only beginning to understand sperm RNA-mediated inheritance in mammals. A recent analysis between obese patients and lean controls identified differences in sperm small RNAs, which could be partially reset in response to aggressive weight loss^5^. Another study identified differential expression of long non-coding RNAs (lncRNAs) between diabetic and non-diabetic mice^6^. Environmental effects caused by traumatic stress induced by MSUS (unpredicted maternal separation combined with unpredictable maternal stress) or chronic stress^7–9^ or a high fat diet during adolescence^10,11^ can be passed down to the next generation through miRNAs or tRNA fragments in sperm. These results suggest a rapid communication between the somatic cells and the germlines, and that the communication leaves reversible marks on sperm RNAs.

Although the mechanisms that determine the sperm RNA profile remain elusive, it has been reported that the tRNA fragments in sperm can be deposited from the epididymis^10,11^, providing one route to control sperm RNA profiles. However, we do not know whether this route affects other RNAs in sperm, such as miRNAs and longer RNAs (>200nt) as it is believed that most sperm RNAs are relics from spermatogenesis. When the meiotic products, round spermatids, package their genomes into compacted nuclei in preparation for delivery to oocytes, a stage known as spermiogenesis, the spermatid genome enters a transcriptionally quiescent state. The majority of cytoplasm in round spermatids is enclosed into cytoplasmic vesicles, the residual bodies, which are engulfed by Sertoli cells and degraded^13^. Ribosomes are removed as a component of the residual body^14^, and only fragmented rRNAs can be detected in sperm^15^. We do not know whether sperm RNAs are randomly left over from spermatogenesis or if there is an active mechanism to select specific RNAs for persistence in sperm.

While the transgenerational function of sperm small RNAs has been established^7,10,11^, we know little about sperm long RNAs. As a consequence of losing 80S ribosomes, total RNA in sperm has a size distribution lacking the 18S and 28S rRNA peaks^16^ (Supplementary Fig. 1a), and is reminiscent of degraded RNA samples. This has led to the suggestion that most long RNAs undergo a massive RNA elimination process during spermiogenesis via an unidentified RNase, leaving degraded products in sperm^15,17^. As cytosolic translation has ceased in sperm, intact long RNAs appear to be not only unnecessary for sperm maturation, but also be a burden for cargo-carrying function of sperm. Because of the notion that sperm long RNAs were most degradation products, sperm RNA analysis has principally been focused on exon fragments (sperm RNA elements, SREs), rather than transcriptional units^18^, and sperm RNA research has been biased towards small RNAs.

Recently, long RNAs in sperm are demonstrated to mediate epigenetic inheritance of certain syndromes induced by MSUS in mice, and the effect is eliminated with fragmented long RNAs^19^. Yet, it is unclear which RNAs exist in their intact form in sperm because until recently, we lacked sensitive, high-throughput experimental techniques that can distinguish intact RNAs from fragmented RNAs. Northern blotting can examine the intactness of mRNAs, but requires significant quantities of input RNA (micrograms), and is low throughput. High-throughput RNA sequencing by Illumina technology (short-read/second-generation sequencing) can only attain short contiguous read lengths (at most 300 bp). Thus, library preparations result in fragmented RNAs, causing transcript integrity loss. Studies based on short-read sequencing have been performed on sperm RNAs^15,18,20,21^, but it has not been established whether and to what extent intact mRNAs exist in sperm.

To test the intactness of sperm long RNA, we will also need precise annotation of transcriptome in testis and sperm. However, the reference transcriptome (Ref-seq) annotation is incomplete and unprecise. Testes, with germ cells (precursors of sperm) constitute ∼92% their cell populations based on the recent single cell sequencing results^22^, usually express specific RNA isoforms that cannot be found in other tissues^23–25^. The short-read length, the loss of linkage information in intron-exon structures, and the uncertain source of the multi-mapping reads lead to a failure in accurately defining transcript structures (e.g., 5’ and 3’ boundaries and splicing diversity), and to a failure in discovery of unannotated transcripts harboring repetitive sequences. Although attempts have been made to assemble transcriptomes based on short reads^26,27^, the lack of information on transcript integrity suggests the problem is unlikely to be overcome computationally, which accounts for another hurdle to identify intact long RNAs in sperm.

Single-molecule long-read sequencing technology (third-generation sequencing; PacBio, Iso-seq, Supplementary Fig. 1b) provides a means to identify full-length transcripts^28^. The PacBio Sequel system can produce long reads (up to 30 kb *vs* 250 bp for short-read sequencing). Although long-read sequencing overcomes many of the problems of short-read sequencing, there are still limitations. The corresponding high error rate in base calling (10-15%) can be reduced by self-error correction, by increasing sequencing depth or circular consensus sequencing/CCS, or by hybrid error correction, using high-quality short reads. Although PacBio Iso-seq yields long-read lengths, determining accurate 5’- and 3’-boundaries of intact transcripts remains challenging. Iso-seq cannot distinguish intact transcripts with an N(5′)ppp(5′)N cap structure (5’-cap) from decay intermediates with a 5’-monophosphate or a 5’-hydroxyl. As revealed in this study, Iso-seq uses oligo-dT primers to capture mRNA polyA tails (Supplementary Fig. 1b), which can produce off-target effects by priming from adenine runs within transcripts (Supplementary Fig. 1c). For example, 17.9% of the long reads in our sequencing libraries result from internal priming. Thus, there is a need for complementary data and computational methods to address these intrinsic problems associated with reconstitution of the transcriptome.

Here, we present a comprehensive characterization of sperm and testis transcriptomes by pairing PacBio Iso-seq sequencing with Illumina sequencing of 5′-ends bearing a 5’cap (cap analysis of gene expression; CAGE^29^) and the 3′-ends preceding the poly(A) tail (polyadenylation site sequencing; PAS-Seq^30^). In total, we identified 2,778 intact transcript species from 1,668 genes in mouse sperm. The SpILT profile is conserved in humans, and they are functionally enriched for protein synthesis functions. The SpILT abundance can be finely tuned by environmental perturbations. In addition to epididymis contribution, most SpILTs and their base-pair complementary sperm miRNAs are coregulated during spermiogenesis through an active post-transcriptionally selection process in both mice and human. In sum, our study reveals that intact long RNAs exist in sperm and identifies conserved selection mechanisms acting to retain certain intact mRNAs in sperm. The identification of the conserved SpILT profile and their abundance changes upon stress expands the potential information carried by sperm, and narrows down the list of RNAs used for diagnostic purposes. Our integrated computational pipeline for identifying intact transcripts and characterizing accurate transcriptomes in sperm and testis provides a valuable resource for studies beyond male reproduction.

## Results

### Acquisition of ultra-pure sperm for RNA preparation

Given that a sperm contains ∼100 fg RNA^31^, and a typical mammalian cell contains 10–30 pg RNA, sperm purification is critical to avoid somatic RNA contamintation^18,32,33^. To collect sperm with high purity, we combined a swim-up procedure with a somatic cell lysis procedure, each of which has been used independently. We collected sperm by letting them swim out of cauda epididymis, and then performed a hypotonic treatment to remove contaminating somatic cells^16,34^. The hypotonic treatment buffer and treatment time were optimized to ensure that sperm integrity was not affected^35^. Using qPCR, we measured the abundance of *Myh11*, a marker for epididymis contamination^36^, to assess the purity of the sperm RNA samples. The presence of *Myh11* in our sperm RNA samples was below the detection limit (<1/10,000, Table S1). Based on these quantification data, we considered our sperm RNA to be ultra-pure.

### Characterization of sperm and testis transcriptomes

We generated 256,879 PacBio Iso-seq long reads to identify the full-length transcripts in purified mouse sperm as described in the paragraph above (Supplementary Table S2), and the depth has reached saturation for isoform identification (Supplementary Fig. 1d). To avoid sequencing-method length biases, we separated the cDNA libraries based on length, and sequenced them separately. We also sequenced the adult mouse testis and analyzed together with sperm sample for two reasons. First, it serves a positive control for PacBio Iso-seq sequencing (Supplementary Fig. 1d). Second, the low yield of sperm RNAs has limits the feasibility to generate CAGE and PAS libraries from sperm. Given that most sperm RNAs are relics from spermatogenesis, sperm RNAs are a subset of testicular RNAs. Thus, CAGE and PAS libraries from testis can be used to correct the pool of intact transcripts from testis and sperm.

Illumina short reads from adult testis and sperm were used to assemble transcripts and correct the single base errors within the PacBio long reads (Fig. 1a, i). The corrected long reads were aligned to the short-read assembled transcripts, and those supported by corrected long reads were used for further analysis (Fig. 1a, ii). Long reads not supported by assembled transcripts were ‘rescued’ (Fig. 1a, iv) as they may represent intact transcripts that the short-read based method may have failed to assemble due to sequence bias or repetitive regions^37^. We used our published high-throughput CAGE data from mouse testes^38^ to identify intact RNAs and refine the transcript 5’-end boundaries (Fig. 1a, iii). Also, we used our published PAS-Seq data from mouse testis^38^ to filter out internally primed artifactual transcripts from the final set of verified intact RNAs (Fig. 1a, v).

**Fig. 1.**
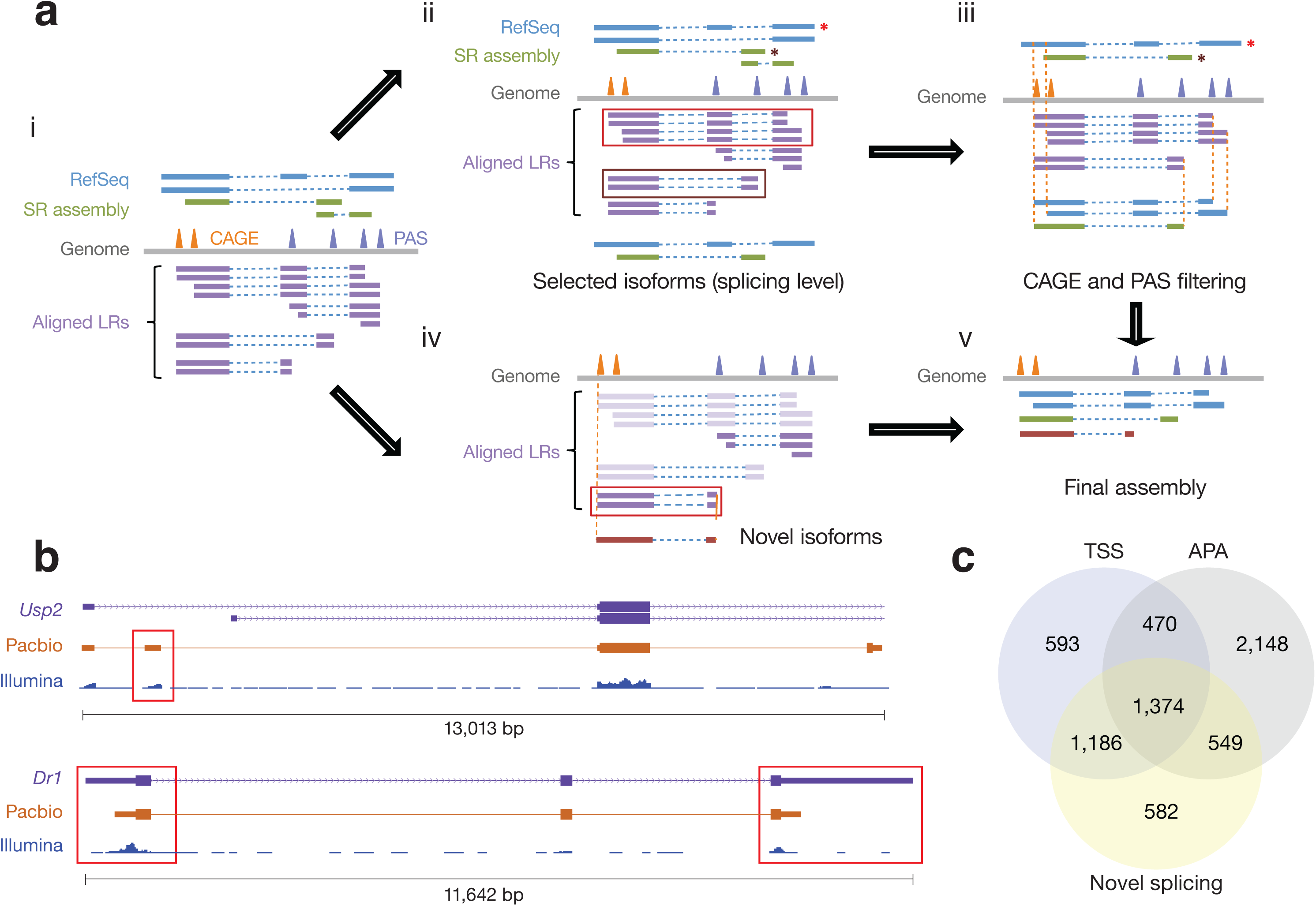
Pipeline for identifying intact transcripts in mouse sperm. a) Strategy to assemble the transcriptome from mouse testis and sperm. The reconstruction of transcriptome mainly contained four steps. **i**, we compared the successfully aligned long reads (LRs) with the Ref-seq annotation of mm10 and assembled the short reads (SRs) using StringTie (version 1.3.1c, parameters -c 10 -p 10)^79^. SRs from sperm and testis were assembled separately and the two assemblies were then merged using StringTie again (the “merge” mode). Peaks from CAGE and PAS-seq were defined by AS-peak^80^. **ii**, Select isoforms that can be supported by any splicing patterns in Ref-seq or SR assembly. It should be noticed that we did not impose any restriction at 5’ or 3’ end at this stage. **iii**, Correct the ends of isoforms by CAGE and PAS peaks. **iv**, Novel isoforms were defined by selecting LRs that are not supported by either Ref-seq or SR assembly. These isoforms must have CAGE and PAS peaks to support their ends. **v**, Final assembly contains well annotated intact RNA in sperm and testis. b) Two examples of novel transcript structures in sperm. From top to bottom, genomic location, Ref-seq, PacBio reads & Illumina reads sequencing from mouse sperm. c) Venn diagrams showing the contribution of alternative transcription start site (TSS), alternative splicing, and alternative polyA sites (APA) to the isoforms that differ from Ref-seq.

Overall, we identified 10,919 intact long transcript species in sperm and testis combined. Only 3,152 transcript species were exactly the same as those deposited in Ref-seq: 865 transcript species were from loci not included in Ref-seq and 6,902 transcript species were novel isoforms from annotated loci (Fig. 1b, Supplementary Fig. 1e). Tissue specific isoforms mainly arose from the use of alternative PolyA sites (APAs), with smaller numbers arising from alternative splicing, or alternative transcription initiation (Fig. 1c). Using computational approaches to scan homolog for sequences and protein domains^39^, we separated these transcript species into 9,388 mRNAs and 1,531 lncRNAs. Taken together, we have generated high-quality reference transcriptomes for mouse sperm and testis.

### Intact long RNAs are present in sperm

Our study reveals the existence of intact long RNA species in sperm. Based on the PacBio Iso-seq data, in sperm we detected 2,778 SpILT species (2,246 mRNAs and 532 lncRNAs) from 1,384 gene loci. To quantify the abundance of SpILTs, we established standard curves for absolute amounts of RNA (using RT-PCR) and for quantifying sperm number (using *in vitro* synthesized DNA). Two SpILT mRNAs, *Rps6* and *Rpl22*, which code for 80S ribosome proteins, were chosen for detection with 82 ± 19 *Rps6* transcripts/sperm and 24 ± 3 *Rpl22* transcripts/sperm (Fig. 2a). Since qPCR amplicons are too short to span the entire transcripts, the qPCR itself could not distinguish decay intermediates from intact transcripts. However, the sequence reads in sperm were evenly distributed throughout the entire ribosomal protein-encoding transcripts, as seen in the RNA-seq in testis (Supplementary Fig. 2a), arguing against the possibility that most of the detected RNAs are decay intermediates. Nevertheless, the quantity we detected rules out the possibility that these long RNAs were retained by tethering to their template DNA^40^. Using the quantification to calibrate the short sequencing results, we estimate around 140,000 SpILTs per sperm, which supports for potential biological function.

**Fig. 2.**
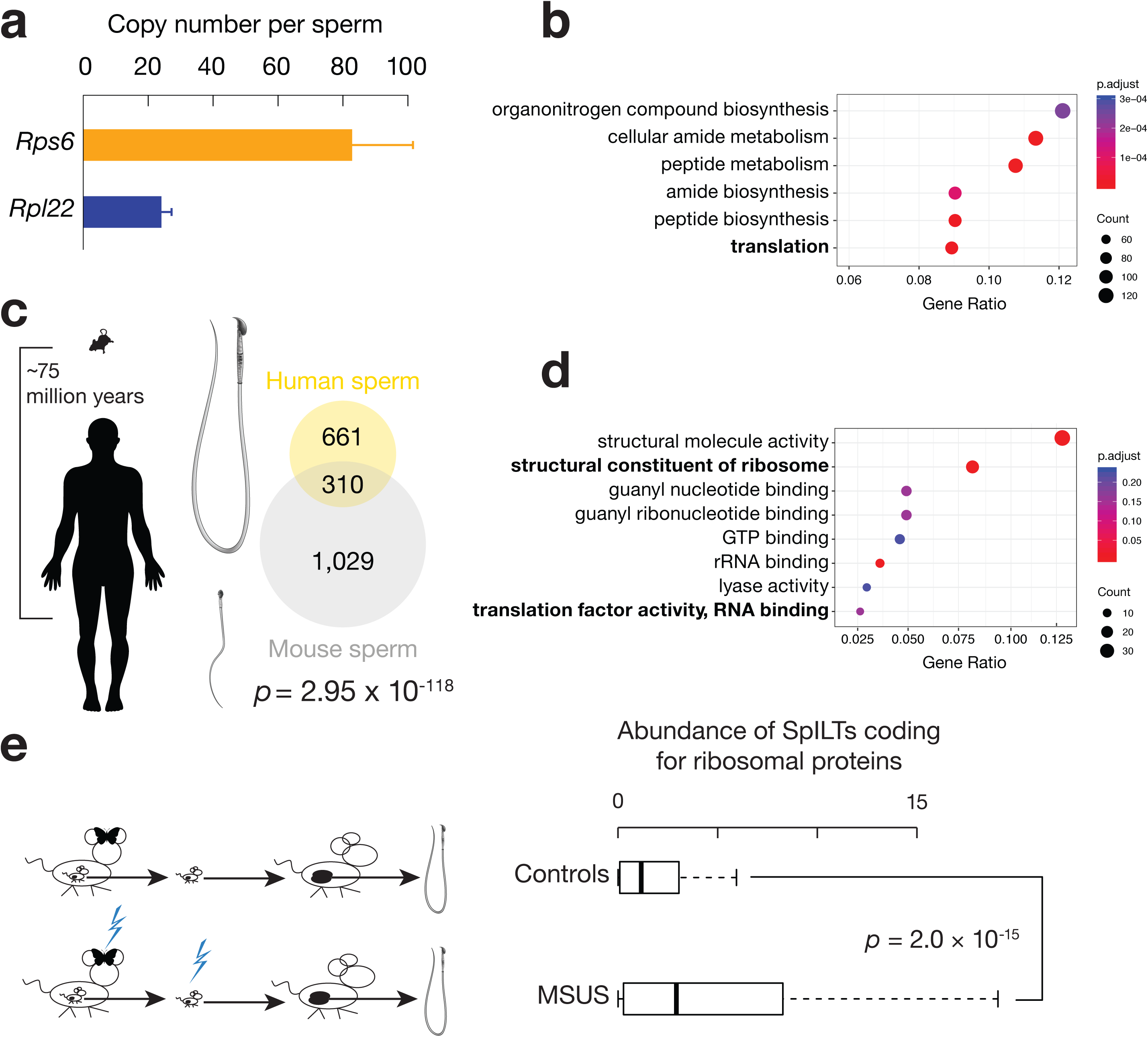
Conserved intact transcripts exist in sperm. a) Copy number per sperm of SpILT mRNAs *Rps6* and *Rpl22*. The RNAs were quantified using *in vitro* transcribed RNAs as standards. Abundance of transcripts was quantified using RT-qPCR (n = 3). Data are mean ± standard deviation. b) Sperm RNAs are enriched for protein synthesis functions. c) Phylogenetic separation of mice and humans as well as the morphology of their sperm. d) Conserved sperm RNAs are enriched for protein synthesis functions. e) Box plot showing abundance of sperm mRNAs coding for ribosomal proteins in control vs MSUS conditions. Paired Wilcoxon Rank Sum and Signed Rank Tests were performed.

### SpILT profiles are evolutionarily conserved

SpILTs functioning in spermatogenesis were not enriched in mice, based on gene ontology analysis (Fig. 2b), unlike the intact transcripts in testis, which is significantly enriched for spermatogenesis function (Supplementary Fig. 2b). This is in contrary to the expectation that SpILTs are random leftovers from spermatogenesis^41–43^. Instead, among the most significant enrichment was in mRNAs encoding the translational apparatuses (Fig. 2b), a function not required during sperm maturation.

To test whether SpILT profiles are evolutionarily conserved, we performed long-read sequencing of RNAs from purified human sperm (Supplementary Fig. 1a), which have morphologically significantly diverged from mouse sperm (Fig. 2c). Among 14,586,344 subreads, a depth that reached saturation for isoform detection (Supplementary Fig. 1d), we detected 1,365 intact mRNAs and 183 lncRNAs. Because of the lack of CAGE and PAS data, we did not consider long reads whose 5’ ends or 3’ ends could not be supported by Ref-seq, and thus the number of intact transcripts in human sperm is underestimated.

We found that 32% (310/971) of human genes coding for SpILTs have mouse homologs that also code for SpILTs, and 23% (310/1339) of mouse SpILT genes also have human SpILT homologs. Of the 17,550 genes shared in both mice and humans, the overlaps between human and mouse SpILT genes were not random (Fig. 2c, Fisher exact test, *p* = 3.0 × 10^−118^). Among these conserved mRNA genes, the genes functioning in spermatogenesis were also not enriched in human sperm, whereas the mRNAs encoding for ribosomal components were enriched (Fig. 2b), as in mouse sperm (Fig. 2d). These results indicated conservation of the SpILT repertoire, which likely predates the divergence of rodents and primates 75 million years ago (Fig. 2c)^44^.

### SpILT abundance in response to environmental perturbations

Long RNAs have been demonstrated to mediate to the inheritance of trauma symptoms induced by MSUS, including food intake, glucose response to insulin and risk-taking in adulthood^19^. Injection of long RNAs, from the sperm of males with MSUS, replicates the specific effects of MSUS in the offspring^19^. To test how SpILT abundance responds to MSUS, we reanalyzed the previously published RNA-seq from sperm^19^. Among 1,384 genes that code for SpILTs in mice, 453 displayed significant changes (*q* < 0.1) with the expression of 416 genes increasing and 37 decreasing (Supplementary Fig. 2c, Table S2). The 453 SpILT were enriched for ribosomal components, based on GO-term analysis (*p* = 3.7 × 10^−37^). The median abundance of ribosomal-encoding SpILTs increased significantly by 2.7 fold (Fig. 2e, *p* = 2.0 × 10^−15^), indicating that the SpILT subclass encoding ribosomal-protein is responsive to mental stresses. Thus, while SpILT profile is stable over evolutionary timescales, their abundance can be fine-tuned by environmental perturbations.

### SpILTs are mostly determined in testis

What regulates SpILT profiles? Considering the transcriptional quiescence of the sperm genome, there are two possible sources of SpILTs: relics from spermatogenesis in testis or deposits from the epididymis. Most of the SpILTs were also detected in testis, supporting the notion that spermatogenesis in testis is their primary origin (Fig. 3a). The transcripts that were found in sperm but not in testis may either be due to their origin in the epididymis or their low abundance in testis. Indeed, we detected long reads from *Crisp1* in sperm but not in testis (Supplementary Fig. 3a). *Crisp1* is expressed in the epididymis, and the protein is secreted into the epididymal lumen, and functions in sperm-egg fusion^45,46^. Our result indicates that *Crisp1* mRNAs were also received by sperm. In additional to depositing tRNA fragments^10,11^, our results indicate that the epididymis indeed contributes to SpILTs.

**Fig. 3.**
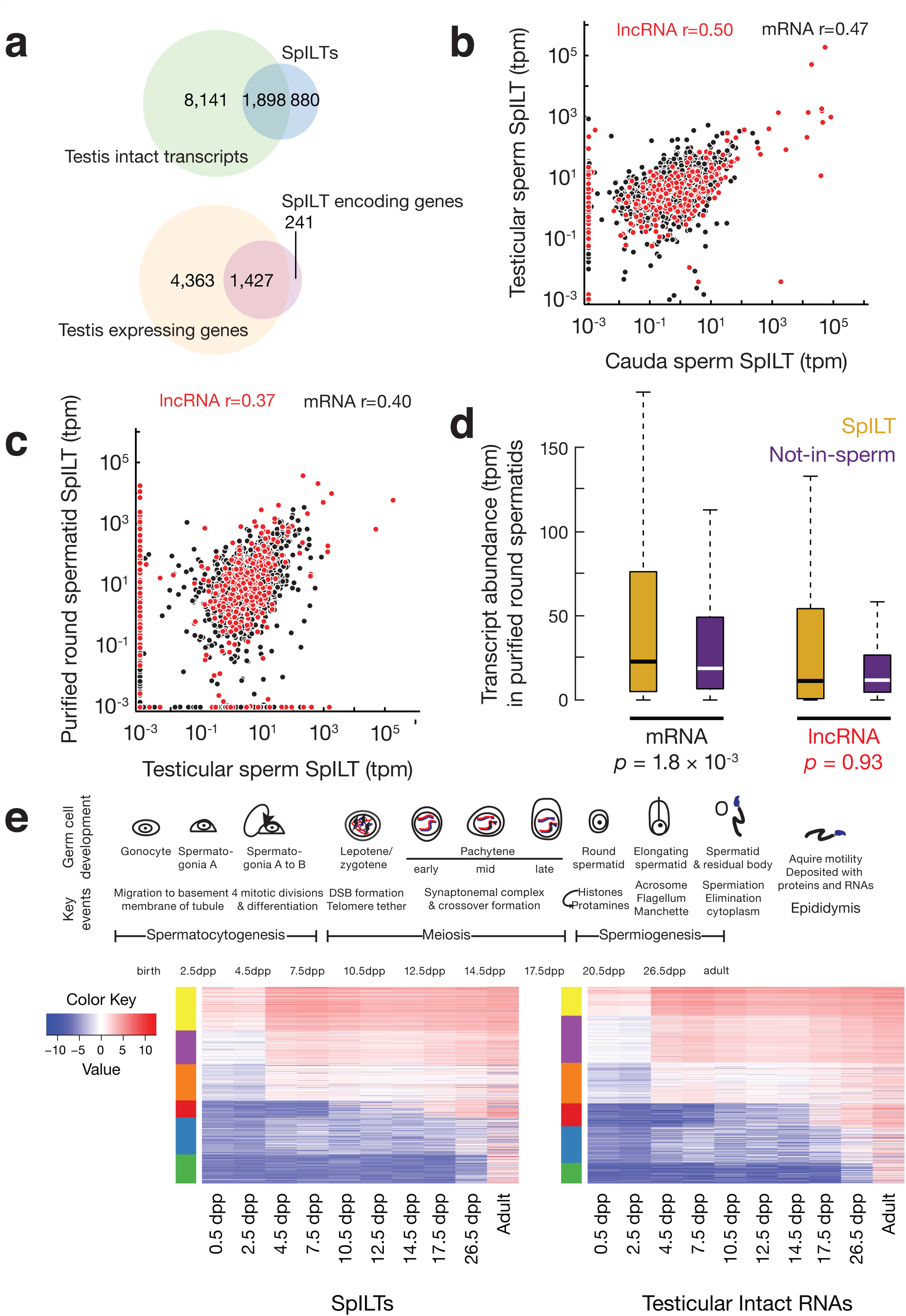
SpILTs are actively selected from round spermatid precursor cells. a) Venn diagrams showing the overlapping intact transcripts (upper) and genes (lower) detected in sperm and in testis. b) SpILT abundance in testicular sperm versus cauda sperm. Tpm, transcripts per million. Pearson correlation coefficient (r) was computed for mRNA and lncRNA separately. c) SpILT abundance in mouse testicular sperm versus mRNA abundance in purified round spermatids. Pearson correlation coefficient (r) was computed for mRNA and lncRNA separately. d) Box plot showing RNA abundance in purified round spermatids. Gold, SpILTs. Purple, intact testicular mRNAs that were not detected in sperm. Wilcoxon Rank Sum and Signed Rank Tests were performed. In 3c and 3d, we analyzed previously published purified round spermatid RNA-seq data^49^. e) Heatmap showing the expression dynamics of testicular RNAs (left) and the expression dynamics of SpILT in mice (right). Key events in mouse spermatogenesis were listed along with the time points. DSB. Double strand break. dpp, days post partum. The RNAs are grouped into 6 clusters (using clara function in R ‘cluster’ package) indicated by colored bars to the left of the heat maps. There was no significant difference in the percentage in each cluster between the SpILTs and testicular intact RNAs (χ^2^ test, *p* = 0.13).

To what extent does the epididymis impact SpILTs? We used high-throughput sequencing of short reads to quantitatively investigate the origins of sperm RNAs. We purified testicular sperm (before their pumping into the epididymis), and compared their RNAs with RNAs from cauda sperm (isolated from the cauda of the epididymis), using spike-in RNAs and sperm count for normalization. We chose young mice that had just reached sexual maturity to avoid the aging effects of long-term sperm storage in the epididymis^47^. Transcript abundance was highly reproducible between biological replicates (Supplementary Fig. 2b). Overall, the SpILT abundance, for either mRNAs or lincRNAs, in testicular sperm and cauda sperm is significantly correlated (Fig. 3b, *p* < 2.2 × 10^−16^). Among the 1,384 genes that code for SpILT in cauda sperm, 35 genes showed significantly higher expression compared to that in testicular sperm (q < 0.1, Table S2), suggesting an epididymal origin. 58 genes showed significantly lower expression (q < 0.1, Table S2) in cauda sperm compared to testicular sperm. These transcripts likely localize in the cytoplasmic droplets that are shed as sperm mature from the testis to cauda^48^. While maturation in epididymis impacts SpILTs, SpILT composition as well as abundance is mostly determined during spermatogenesis in testis.

### Neither RNA abundance nor transcription dynamics in testis distinguish SpILTs

Transcripts may persist in sperm either through random acquisition from precursor cells or by an active selection mechanism operating during sperm formation. To distinguish between these hypotheses, we analyzed correlations between RNA abundance in sperm and their abundance in precursor cells. Considering that transcription ceases after the round spermatid stage (which is followed by subsequent morphology changes), we used purified round spermatids as the closest precursors of mature sperm^49^. The correlation between SpILT abundance in purified round spermatids and abundance in testicular sperm is weak to support the random acquisition hypothesis (Fig. 3c). The abundance distribution of lncRNAs that are, or are not, found in sperm displayed no obvious difference in purified round spermatids (Fig. 3d, right, *p* = 0.93). In purified round spermatids, the median abundance of mRNAs that end up in sperm is only 1.2 fold of the abundance of mRNAs that do not end up in sperm (Fig. 3d). These findings on RNA abundance (Fig. 3c) and correlation (Fig. 3d) are consistent with findings using the round spermatid datasets from an independent laboratory (Supplementary Fig. 3c&d)^50^. It is thus likely that other factors determine RNA transcript abundance in sperm.

Although transcription is believed to shut down after the round spermatid stage, it is possible that some transcription may continue and that newly transcribed transcripts end up in sperm. Given the difficulty of purifying the elongating spermatids, we tested this possibility by examining expression dynamics during spermatogenesis. We sequenced total testicular RNAs at 10 time points after birth from wild-type C57BL/6 mice. Since the first wave of spermatogenesis in mice is synchronized, RNA expression in prepubertal testes^51^ can follow spermatogenesis (Fig. 3e). Overall, transcription dynamics did not distinguish the subset of testicular transcripts retained in sperm from those that were eliminated (χ^2^= 0.13, Fig. 3e), arguing against the possibility that SpILTs are determined by transcriptional regulation during late spermiogenesis. Furthermore, SpILT chromatin regions had comparable accessibility to the chromatin regions coding for testicular RNAs not found in sperm, as determined by assay for transposase-accessible chromatin using sequencing (ATAC-seq) in sperm^52^ (Supplementary Fig. 3e), further arguing against the possibility that SpILTs are transcribed during late spermiogenesis. In summary, neither the transcript abundance nor the transcription dynamics distinguishes the SpILTs from the rest of the testicular transcripts, rejecting the random acquisition hypothesis.

### Active post-transcriptional selection of RNAs to retain in sperm

SpILT mRNAs are significantly shorter, less than three quarter of the length of testicular mRNAs that are not found in mouse sperm (Fig. 4a, *p* < 2.2 × 10^−16^). To test the impact of transcript length in sperm RNA retention, we performed a logistic regression that predicts whether an mRNA can be found in sperm given: (1) its expression levels in spermatids, and (2) the log of its lengths. For mRNAs, length is a significant predictor of whether they can be found in sperm (*p* < 2.0 × 10^−16^): when the length is doubled, the chance that it can be found in sperm is reduced by 60.1% (95% CI: 57.5% ∼ 62.5%), while adjusting for the expression level in spermatids. The preference for shorter transcripts is also seen for lncRNAs (Fig. 6a, *p* = 2.9 × 10^−9^), whose length is also a significant predictor of occurrence in sperm (*p* < 1.0 × 10^−8^): Doubling length reduces occurrence by 37.8% (95% CI: 26.9% ∼ 47.2%), while adjusting for expression level in spermatids. Therefore, the fact that shorter mRNAs and lncRNAs preferentially end up in sperm demonstrates that SpILT is controlled post-transcriptionally during spermiogenesis.

**Fig. 4.**
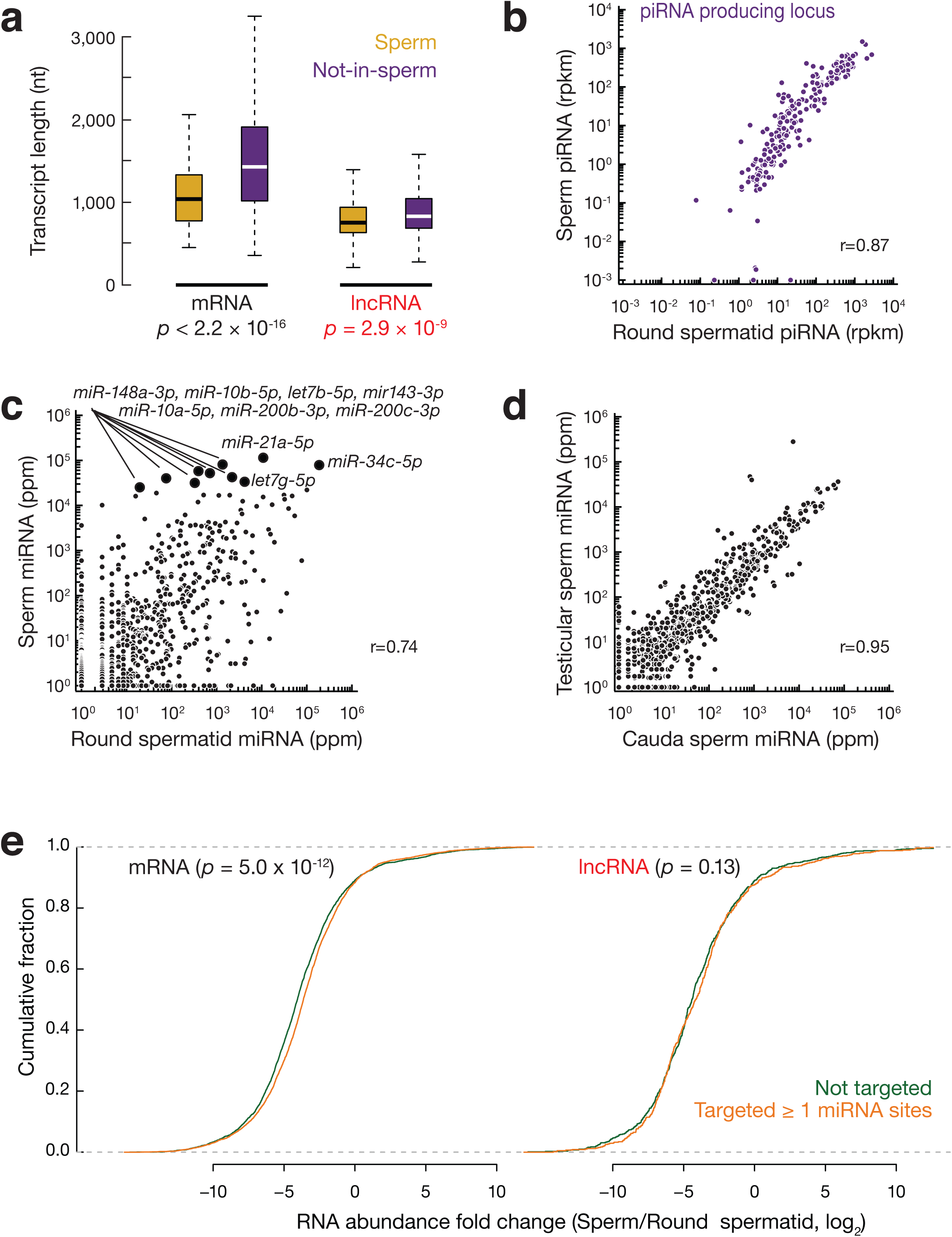
SpILTs and sperm miRNAs are co-regulated. a) Box plot showing transcript lengths. Gold, SpILTs. Purple, intact testicular mRNAs that were not detected in sperm. Wilcoxon Rank Sum and Signed Rank Tests were performed. b) A scatterplot of piRNA abundance from each piRNA producing locus in sperm vs. in round spermatids in mice. rpkm, reads per kilobase pair per million reads mapped to the genome. Pearson correlation coefficient (r) was computed. c) A scatterplot of miRNA abundance in sperm vs. in round spermatids in mice. The top 10 most abundant miRNAs in sperm were marked with giant black circles. Ppm, parts per million. Pearson correlation coefficient (r) was computed. d) A scatterplot of miRNA abundance in testicular sperm vs. in cauda sperm in mice. Pearson correlation coefficient (r) was computed. e) miRNA binding sites are enriched within mRNAs that retained in sperm. Plotted were cumulative distributions of RNA fold changes in sperm to those in round spermatids, left, mRNAs, right, lncRNAs. We used top 10 most abundant miRNAs detected in sperm small RNA-seq and used MiRanda for the target scan^81^.

To test whether transcript length influences the sperm SpILT population in human, we performed a partial correlation analysis^53^ between sperm mRNA abundance and transcript length (while controlling for mRNA abundance in purified round spermatids^49^), given the lack of data on intact transcripts in health adult human testis. We found that mRNA and lncRNA abundances in sperm displayed significant negative correlations with transcript length (mRNA, r = −0.08, *p* = 2.7 × 10^−3^; lncRNA, r = −0.27, *p* = 2.1 × 10^−4^). The preference for short transcripts in both mice and human explains a relatively shorter bioanalyzer total RNA profile of sperm (Supplementary Fig. 1a). In summary, rather than transferring from sperm precursors to sperm randomly, we reveal an active selection mechanism controlling SpILT profiles that are evolutionarily conserved between mice and humans.

### Sperm miRNAs are also mostly determined in testis

Given the known transgenerational effects of sperm small RNAs^7,10,11^, we tested whether sperm small RNAs are also regulated post-transcriptionally during spermiogenesis. Two small-RNA (18–45 nt) sequencing libraries were prepared from sperm: one comprising all small RNAs and one selected for 2’-*O*-methyl-modified 3’ termini, found in PIWI-interacting RNAs (piRNAs), but not in miRNAs. Small-RNA sequencing detected abundant miRNAs and piRNAs in sperm (Supplementary Fig. 4a). Compared to round spermatids, there is a general decrease of piRNA abundance in sperm, but the abundance of piRNAs, either mapping to individual piRNA producing loci or individual transposon families, are highly correlated between round spermatids and sperm (Fig. 4b, Supplementary Fig. 4b). In contrary to piRNAs, the abundance of sperm miRNAs poorly correlates with that in round spermatids (Fig. 4c). The weak correlation is not due to the sperm maturation outside of testis, as the miRNA abundance between testicular sperm and cauda sperm is highly correlated (Fig. 4d, Supplementary Fig. 4c&d). Thus, similar to SpILTs, a specific selection mechanism during spermiogenesis determines sperm miRNA profiles.

### miRNAs and SpILT mRNAs are co-regulated

We found that SpILT mRNAs harbor more miRNA binding sites compared to testicular mRNAs that are not retained in both mice and human sperm. We selected the 10 most abundant miRNAs in mouse sperm (Fig. 4c), used their seed sequences to predict targets site on mRNAs, and plotted their abundance ratio in sperm to that in round spermatids. We found that miRNA binding sites are enriched within SpILT mRNAs in mice (one-sided Kolmogorov–Smirnov [K–S] test, *p* values = 1.1 × 10^−16^, Fig. 4e, left). No such enrichment was seen in SpILT lncRNAs (*p* values = 0.59, Fig. 4e, right), consistent with the notion that miRNAs mostly target mRNAs^54^. A similar enrichment of miRNA binding sites of the 10 most abundant miRNAs in human sperm was observed for SpILT mRNAs, and not lncRNAs, in human (Supplementary Fig. 4e). Given that shorter mRNAs are preferentially found in sperm, the higher number of miRNA binding sites on SpILT mRNAs is not simply due to transcript length. Our results imply that base-complementary miRNAs and mRNAs are retained in sperm together.

Given that sperm small RNAs and long RNAs function synergistically to replicate the symptoms of MSUS in the offspring^19^, we next tested whether miRNAs and their mRNA targets are co-regulated in MSUS. Five miRNAs—miR-375-3p, miR-375-5p, miR-200b-3p, miR-672-5p and miR-466-5p—have been reported to increase in abundance in F1 MSUS sperm^7^. We performed a target search of these miRNAs on SpILT mRNAs and found that the target mRNAs also increased in abundance in MSUS compared to controls (Supplementary Fig. 4f, left). Such enrichment was not observed for SpILT lncRNAs in MSUS (Supplementary Fig. 4f, right). miRNA binding could be a driving force for SpILT mRNA selection. Alternatively, if miRNA-independent mechanisms retain mRNAs in sperm, the mRNAs could function as “sponges” to retain miRNAs. Nevertheless, our results indicate that in both mice and human, both under normal physiological conditions and during environmental perturbations, SpILT mRNAs are co-regulated with their base-complementary miRNAs.

## Discussion

In this study, we demonstrate that intact long RNAs are found in sperm. SpILT profiles are evolutionarily conserved between rodents and primates. SpILT mRNAs are enriched for protein synthesis functions, the exact functional subsets of which is sensitive to mental stress. SpILTs found in sperm comprise a subset of RNAs found in precursor cells, with transcript length biased with selection. The retention of SpILTs are co-regulated with miRNA retention in sperm.

We have developed strategies and standards for defining intact transcripts. We integrated PacBio Iso-seq long reads with various second-generation sequencing techniques (Fig. 1a): (1) Illumina RNA sequencing was used to correct error-prone long reads and assist in transcript assembly; (2) CAGE sequencing was used to identify the 5’-ends of intact transcripts; and, (3) PAS sequencing was used to identify artifactual alternative PolyA sites due to internal oligo-dT priming. We defined 10,919 intact transcripts in sperm and testis, considerably less than the > 60,000 transcripts identified by short-read assembly, further demonstrating the efficacy of our transcriptome reconstitution pipeline. Only a quarter of the identified transcripts were found in Ref-seq. The majority of isoforms from known genes arose from APA, consistent with observations that proximal PolyA sites are preferred during spermatogenesis^55–57^. The hybrid sequencing and analysis approach used here can be applied to other tissues and organisms.

The identification of SpILTs may shed light on the roles that sperm RNAs may play in the offspring. There are no intact 80S ribosomes in sperm, thus SpILT RNAs are unlikely to be functional until they meet oocyte ribosomes after fertilization. The transgenerational role of SpILTs is supported by the loss of transgenerational effects of long RNAs in MSUS sperm after *in vitro* RNA fragmentation^19^. Given that protein synthesis is activated shortly after fertilization^58,59^, sperm mRNAs that encode ribosomal proteins may contribute to quicker and more robust protein synthesis that can direct more resources from the mother to support the pregnancy. The mental stress from MSUS may predispose their offspring to further enhance the tendency. This idea is further supported by data showing that treating sperm with RNase reduces the body weight of the offspring^60^.

It remains unclear how environmental signals establish epigenetic changes in the germline. Germ cells are sequestered from somatic cells early in development, and sperm production is protected in a specialized niche. This “Weismann barrier” hypothesis provides a strict demarcation between soma and germ lines, restricting the flow of hereditary information from germline to soma and preventing hereditary information flow in the opposite direction^61^. This paradigm is challenged by the identification of transgenerational effects, which indicate routes for soma-to-germline communication. We found that SpILTs are mostly determined post-transcriptionally during spermiogenesis. We assessed whether SpILTs accumulate randomly or are enriched by an active selection mechanism. We eliminate the random accumulation hypothesis by showing that: (1) spermatogenesis GO functions are not enriched in sperm; (2) RNA abundance in round spermatids has only a weak correlation with RNA abundance in testicular sperm; and (3) transcript length contributes to selection. Therefore, an active selection mechanism controls SpILT composition during spermiogenesis. The existence of a selection mechanism for SpILTs may provide a rapid route for external changes to act on sperm RNA plasticity, independent of the epididymis-mediated communication between soma and germline^10,62^.

## Methods

### Animals

C57BL/6J (Jackson Labs, Bar Harbor, ME, USA; stock number 664) mice were maintained and used according to NIH guidelines for animal care and the University Committee on Animal Resources at the University of Rochester.

### Mouse sperm purification

After dissecting adult mice, cauda epididymides were excised and placed into 1 ml Whitten’s-HEPES buffer pH=7.3 (100 mM NaCl, 4.4 mM KCl, 1.2 mM KH2PO4, 1.2 mM MgSO4, 5.5 mM Glucose, 0.8 mM Pyruvic acid, 4.8 mM Lactic acid (hemicalcium), and HEPES 20 mM) at 37°C. Two additional cuts were made on each epididymis to release sperm. After incubation for 20 minutes, the sperm suspension was carefully transferred into a 1.5 ml Eppendorf tube, avoiding as much as possible collection of any contaminating cauda tissue. The sperm sample was spun down at 5000 rpm, rt, 3 min, washed with ddH_2_O, and spun down again. To further eliminate somatic cell contamination, the pellet was re-suspended in 500 μl somatic cell lysis buffer (0.1% SDS, 0.05% Triton-X100) at room temperature. After 5 minutes, precipitate the sperm by centrifuging at 10,000 rpm, room temperature for 3 minutes. Finally, the liquid was completely removed, and the pellet was ready for sperm RNA purification.

Testicular sperm was purified, as previously described^10^, and the purity was verified by microscopic examination. Stock Isotonic Percoll (SIP) was made by mixing Percoll (GE Healthcare, Catalog 17-0891-01) with 1.5 M NaCl solution (9:1). For use, 52% isotonic Percoll was diluted with 0.15 M NaCl on ice. Then 10.5 ml solution was loaded into an ultracentrifuge tube (Beckman, Catalog 331374). Two mouse testes were freshly excised and immediately minced in a 35 mm petri dish containing 1 ml 0.9% NaCl. The suspension was filtered using a cell strainer (Falcon, Catalog 352360), and loaded into the tube containing 52% isotonic Percoll. The tube was centrifuged at 15,000 rpm for 10 minutes at 10°C. The pellet (< 0.5 ml) containing red blood cells, Sertoli cells, and sperm was transferred into a 1.5 ml tube followed by addition of 0.9% NaCl. The tube was centrifuged at 1,500 rpm at 4°C for 10 minutes, and the supernatant was discarded carefully. To lyse the somatic cells, 0.5 ml lysis buffer (0.01% SDS and 0.005% Triton X-100 in water) was added to the pellet. After gently resuspending the pellet, the suspension was incubated on ice for 10 minutes. Finally, the tube was centrifuged at 3,000 g for 5 minutes at room temperature. The pellet containing purified testicular sperm was used directly in experiments.

### Human sperm purification

For conducting these experiments, de-identified normozoospermic (>15 million spermatozoa/ml and >40% motility) semen samples discarded after routine semen analysis were obtained from the Fertility Clinic at the University of Rochester, NY. The study was approved by University of Rochester Medical Center Review Board Protocol 00003599. Any existing identification labels were removed and samples were coded. Sperm were prepared using a 90% PureCeption gradient (Sage Biopharma a Cooper Surgical Company, Trumbull, CT). One to two ml of 90% gradient was added to a 15-ml sterile conical tube (Corning), and 1.5–2 ml of semen sample was layered gently on top of the gradient using a sterile transfer pipette. If semen volume was more than 2 ml, we used more than one tube. The sample was centrifuged at 500 × g for 20 minutes. After centrifugation seminal plasma and 90% gradient was carefully removed using a transfer pipette leaving a small volume of gradient along with the sperm pellet at the bottom. The sperm pellet was transferred to another clean 15 ml tube, and 5 ml of Quinn’s Sperm washing buffer (Hepes-HTF medium with 5 mg/ml human serum albumin) was added, the pellet was resuspended, and centrifuged for 7 minutes. The sperm pellet was resuspended in 1 ml Sperm wash buffer, and cryopreserved using commercially available Arctic Sperm Cryopreservation Medium (Irvine Scientific, Santa Ana, CA). This medium contains a mixture of glycerol and sucrose with serum in an isotonic salt solution, and is used commonly in Fertility labs/Sperm bank. Cryopreservation solution (0.33 ml) was added dropwise to the tube containing 1 ml of sperm suspension at room temperature, and mixed gently to obtain (3:1) mixture of sperm and cryopreservation solution. This mixture was aliquoted in 2-3 cryovials (Corning, NY), secured on an aluminum cryocane, immediately snap frozen in the vapor phase of liquid nitrogen for 30 minutes, and then plunged into the liquid nitrogen. The coded vials were stored in liquid nitrogen storage tanks specifically designated for research samples in the lab until used.

### Sperm RNA purification

Before RNA purification, the sperm pellet was lysed in 100 µl lysis buffer (1% SDS, 0.1M DTT) at 37°C for 10 minutes. After incubation, 300 µl Trizol (Thermo Fisher Scientific, Waltham, MA, Catalog #15596018) was added, and the solution was vortexed for 15 seconds, and then placed on an Eppendorf Thermomixer 5436 (Brinkmann Instruments, Inc. Westbury, New York) and vortexed for another 5 minutes at room temperature. Chloroform (60 µl) was added, and the solution was vortexed for 15 seconds, then placed on an Eppendorf Thermomixer 5436 and vortexed for another 3 minutes at room temperature. The sample was then centrifuged (12,000 g, 4°C, 15 minutes), and the liquid phase was transferred to a clean tube. To precipitate RNAs, an equal amount of isopropyl alcohol 2 µl glycogen was added to the solution. After incubation at room temperature for 10 minutes, the precipitated RNA was pelleted by centrifugation (12,000 g, 4°C, 10 minutes). After an additional wash with 75% ethanol, the pellet was dried at room temperature for 5 minutes. Finally, the RNA was dissolved in 43 µl ddH_2_O.

To eliminate DNA contamination in the RNA samples, 5 µl of 10X Turbo DNase buffer and 2 µl Turbo DNase (Thermo Fisher Scientific, Waltham, MA, Catalog AM2238) were added to the 43 µl of purified RNA. After incubation at 37°C for 30 minutes, 200 µl ddH_2_O and 250 Ml Acid Phenol: Chloroform (Thermo Fisher Scientific, Waltham, MA, Catalog AM9722) was added, and the sample was mixed thoroughly before centrifugation at 12,000 g at room temperature for 15 minutes. After centrifugation, the RNA was precipitated by adding 3 volumes of 100% ethanol to 1 volume of supernatant, and 2 µl of glycogen. After a 1 hour incubation on ice, the precipitate was pelleted at 15,000 g, 4°C, 30 minutes. After an additional wash with 75% ethanol, the pellet was dried at room temperature for 5 minutes. Finally, the RNA was dissolved in ddH_2_O for further library construction.

### PacBio Iso-Seq™ Library Construction and Sequencing

Full-length RNA sequencing libraries (i.e., Iso-Seq™) were constructed according to the recommended protocol by PacBio (PN 101-070-200 Version 05 (November 2017)^63,64^) with a few modifications. Briefly, for analysis of testis, mRNA were assessed on the Agilent BioAnalyzer or TapeStation, and only preparations with a RIN≥ 8 were used for sequence analysis. No such RIN requirement was imposed for analysis of sperm RNA. If needed, the ZYMO Research RNA Clean and Concentrator kit (Cat. # R1015) was used for concentrating dilute samples. RNA preparations considered suitable for IsoSeq typically displayed absorbance ratios of A260/280 = 1.8-2.0 and A260/230 > 2.5.

RNA preparations of similar quality from adult mouse testis and sperm were used for constructing PacBio IsoSeq libraries (SMRT bell libraries). Full-length cDNA was synthesized using the Clontech SMARTer PCR cDNA Synthesis kit (Cat. # 634925) (Clontech, Palo Alto, CA). Approximately 13-15 PCR cycles were required to generate 10-15 µg of ds-cDNA from a 1 µg RNA sample. The library construction steps included: ExoVII treatment, DNA Damage Repair, End Repair, Blunt-end ligation of SMRT bell adaptors, and ExoIII/ExoVII treatment. This procedure resulted in 1.5 microgram of SMRT bell library (i.e., 25-30% yield when starting with 5 microgram of full-length cDNA). Full-length total cDNA was size selected on the ELF SageSciences system (Electrophoretic Lateral Fractionation System) (SageScience), using 0.75% Agarose (Native) Gel Cassettes v2 (Cat# ELD7510), specified for 0.8-18 kb fragments. Fragments were collected in two size ranges, short (0.6-2.5 kb) and long (>2.5 kb). Generation of >2.5 kb fragments required 4 more cycles of amplification and further cleaning. Short and long fragments were combined in an equimolar pool.

For mouse sperm and testis libraries prepared using the PacBIo RSII platform, we collected 12 cDNA fractions of various sizes between 0.8 and ∼15 kb. These fractions were pooled in such a way as to generate cDNA pools over four size ranges (0.8-2 kb, 2-3 kb, 3-5 kb, and >5 kb) for every sample. Further amplification was needed to generate enough material (for library construction) for the two larger size bins. Additional amplification of the larger size bins resulted in small size byproducts. Therefore, a second size selection (for 3-5 and >5 kb fragments) was performed using an 11×14 cm agarose slab gel. Library-polymerase binding was done at 0.01-0.04 nM (depending on library insert size) for sequencing on the PacBio RSII instrument. Diffusion loading was used for the short fragments, and MagBead loading was used for the larger fragments.

Sample cleaning of SageELF fractions and throughout SMRT bell library construction was done following the manufacturer’s protocol (PACBIO). In brief, fractions were purified using AMPure magnetic beads (0.6:1.0 beads to sample ratio). Final libraries were eluted in 15 µL of 10 mM Tris HCl, pH 8.0. Library fragment size was estimated by the Agilent TapeStation (genomic DNA tapes), and these data were used for calculating molar concentrations. Between 75-125 picomoles of library from each size fraction were loaded onto two SMRT cells for PacBio RS II sequencing. Between 5-8 picomoles of library were loaded (diffusion loading) onto the PacBio SEQUEL sample plate for sequencing. For optimal read length and output, libraries were sequenced on LR SMRT cell, using 2 hr pre-extension, 20 hr movies and 2.1 chemistry reagents (for binding and sequencing). All other steps in sequencing were done according to the recommended protocol by the PacBio sequencing calculator and the *RS Remote Online Help* system. For PacBio RS II, one SMRT cell run generated 60-70 thousand reads with a polymerase read length of 13-16 kb. For PacBio SEQUEL, one SMRT cell run generated 500-70,000 reads with an average polymerase read length of ∼27 kb.

### Illumina RNA sequencing

Strand-specific RNA-seq libraries were constructed following the TruSeq RNA sample preparation protocol as previously described with modifications^38,65^. Spike-in RNA, synthetic firefly luciferase RNA (10^−6^ ng per million sperm) was added right before RNA extraction to normalize RNAs in different libraries. We treated the total sperm RNAs with Dnase first, and then performed the rRNA-depletion with complementary DNA oligos (IDT) and RNaseH (Invitrogen, Waltham, MA, USA)^66,67^. Single-end sequencing (126 nt) of libraries was performed on a HiSeq 2000.

### Small RNA sequencing library construction

Small RNA libraries were constructed and sequenced as previously described^68,69^ using oxidation to enrich for piRNAs by virtue of their 2’-*O*-methyl-modified 3’ termini^69^. A 25-mer RNA with 2’-*O*-methyl-modified 3’ termini (5’-UUACAUAAGAUAUGAACGGAGCCCmG -3’) was used as a spike-in control (5 × 10^−3^ ng per million sperm) according to sperm counts before total RNA extraction. Single-end, 50 nt sequencing was performed using a HiSeq 2000 instrument (Illumina, San Diego, CA, USA).

### Quantitative reverse transcription PCR (qRT–PCR)

Extracted RNAs were treated with Turbo DNase (Thermo Fisher, Waltham, MA, USA) for 20 min at 37°C, and were then size-selected to isolate RNA ≥ 200 nt (DNA Clean & Concentrator™-5, Zymo Research) before reverse transcription using All-in-One cDNA Synthesis SuperMix (Bimake, Houston, TX, USA). Quantitative PCR (qPCR) was performed using the ABI Real-Time PCR Detection System with SYBR Green qPCR Master Mix (Bimake). Data were analyzed using DART-PCR^70^. Spike-in RNA was used to normalize RNAs in different samples. Table S2 lists the qPCR primers.

### Transcriptome reconstruction and intact mRNA identification

The PacBio long reads (LRs) were extracted from the bax files using SMRT Analysis pipeline (version 2.3.0). Isoseq-classify (default parameters) was then used to identify non-chimeric long reads in which all the 5’ primer sequences, polyA ends, and 3’ primer sequences were present. Those long reads which did not satisfy these conditions were filtered out. LoRDEC (version 0.5.3, parameters -k 17 -s 3 -a 50000) was used to perform error correction on the LRs with SRs as input. The corrected LRs were aligned to the mm10 reference genome using GMAP (version 2014-12-24, parameters -t 10 -B 5 -A --nofails -f samse -n1). The aligned LRs with clipping regions longer than 100-nt at either end were filtered out. The SRs were aligned to the mm10 reference genome using HISAT2 (version 2.0.4, parameters --ignore-quals --dta -- threads 10 --max-intronlen 150000).

The reconstruction of transcriptome consisted of four main steps (Fig. 1a). First, we compared the successfully aligned LRs with the RefSeq annotation of mm10, and recorded the isoforms that had full-length LRs. It should be noticed that we did not impose any restriction at the 5’ or 3’ end at this stage. In the second step, we assembled the SRs using stringtie (version 1.3.1c, parameters -c 10 -p 10). SRs from sperm and testis were assembled separately, and the two assemblies were then merged using stringtie again (the “merge” mode). The successfully aligned LRs were further compared with the merged SR assembly, and the isoforms which had full-length LRs were recorded. In the third step, we selected the LRs that were supported by CAGE peaks at 5’ ends. We performed cluster on the CAGE supported LRs that were not identified as full-length LRs in the above two steps; this process identified a set of novel isoforms. To reduce false positives, we only retained clusters that contained at least 2 LRs. The novel isoforms were combined with those recorded in the above two steps. In the last step, we inferred the TSS and polyA isoforms at 5’ and 3’ ends. Specifically, we collected the CAGE supported full-length LRs with respect to each isoform. Each collected LR corresponded to a pair of coordinates marking the positions of the CAGE peak and the 3’ end of the LR on the reference genome. By clustering the coordinate pairs, we obtained a detailed annotation of each TSS-polyA isoform.

The identification of intact mRNAs was completed along with the transcriptome reconstruction. Intact mRNAs were the LRs whose 5’ ends were supported by the CAGE peaks or known Ref-seq transcripts, and whose 3’ ends were supported by the PAS reads or known Ref-seq transcripts.

### General bioinformatics analyses for Illumina sequencing

Analyses were performed using piPipes v1.4^71^. All data from the RNA sequencing, CAGE, and PAS sequencing were analyzed using the latest mouse genome release mm10 (GCA_000001635.7) and human genome release hg38 (GCA_000001405.27). Generally, one mismatch was allowed for genome mapping. Relevant statistics pertaining to the high-throughput sequencing libraries constructed for this study are reported in Table S2.

For small RNA sequencing, libraries were normalized to the sum of total miRNA reads; spike-in RNA was used to normalize the different libraries. Pre-miRNA and mature miRNA annotations were taken from miRbase 22 release^72^. 214 piRNA precursors defined in our previous studies with mm9^38^ were converted to mm10 coordinates with using *liftOver*^73^ with minor manual correction. We used 1,223 mouse transposon families that were defined in both Repeat Masker from UCSC^74^ and Repbase (Jurka et al., 2005). We used 308 mouse tRNAs including 22 mitochondrial tRNAs downloaded from UCSC and Mouse Genome Informatics^75^ and 286 nuclear tRNAs downloaded from UCSC. tRNAs with the same sequences are combined into one species. Uniquely mapping reads >23 nt were selected for piRNA analysis. Oxidized sample from testis was calibrated to the total small RNA samples in the corresponding genotypes via the abundance of shared, uniquely mapped piRNA species. We analyzed previously published small RNA libraries from round spermatids from adult wild-type mouse testis (GSM1096604)^38^ and human sperm (GSM1375214, GSM1375215)^76^. We report piRNA abundance either as parts per million reads mapped to the genome (ppm), or as reads per kilobase pair per million reads mapped to the genome (rpkm) using a pseudo count of 0.001. We report miRNA abundance either as parts per million reads mapped to the genome (ppm) using a pseudo count of 1.

For RNA-seq reads, the expression per transcript was normalized to the top quartile of expressed transcripts per library calculated by Cuffdiff^26^ or to the spike in RNAs, and the tpm (transcript per million) value was quantified using the Salmon algorithm^77^ using a pseudo count of 0.001. We analyzed our previously published RNA-seq library from C57BL/6J wild-type mouse testis at 10.5 dpp (GSM1088421), 12.5 dpp (GSM1088422), 14.5 dpp (GSM1088423), 17.5 dpp (GSM1088424), 20.5 dpp (GSM1088425) and adult stage (GSM1088420)^38^, round spermatids from C57BL/6 control mice at 26dpp (GSM1561107, GSM1561108, GSM1561109, GSM1561110)^50^, round spermatids from adult CD1 mice (GSM1674008 and GSM1674021)^49^, and sperm from MSUS mice (ERR2012004, ERR2012005, ERR2012006, ERR2012007, ERR2012008, ERR2012009, ERR2012010) and control mice (ERR2012000, ERR2012001, ERR2012002, ERR2012003)^19^.

PAS-seq libraries were analyzed as previously described^38^. We first removed adaptors and performed quality control using Flexbar 2.2 (http://sourceforge.net/projects/theflexibleadap) with the parameters “-at 3 -ao 10 --min-readlength 30 --max-uncalled 70 --phred-pre-trim 10.” For reads beginning with GGG including (NGG, NNG or GNG) and ending with three or more adenosines, we removed the first three nucleotides and mapped the remaining sequence with and without the tailing adenosines to the mouse genome using TopHat 2.0.4. We retained only those reads that could be mapped to the genome without the trailing adenosine residues. Genome-mapping reads containing trailing adenosines were regarded as potentially originating from internal priming, and thus were discarded. The 3′ end of the mapped, retained read was reported as the site of cleavage and polyadenylation. We included our previously published PAS-seq library from wild-type adult mouse testis (GSM1096581) in this analysis^38^.

CAGE was analyzed as previously described^38^. After removing adaptor sequences and checking read quality using Flexbar 2.2 with the parameters of “-at 3 -ao 10 --min-readlength 20 --max-uncalled 70 --phred-pre-trim 10,” we retained only reads beginning with NG or GG (the last two nucleotides on the 5′ adaptor). We then removed the first two nucleotides, and mapped the sequences to the mouse genome using TopHat 2.0.4. The 5′ end of the mapped position was reported as the transcription start site. We included our previously published CAGE library from wild-type adult mouse testis (GSM1096580) in this analysis^38^.

ATAC-seq were analyzed using ENCODE ATACseq pipeline (https://www.encodeproject.org/atac-seq/). After genome mapping using bowtie2 and peak calling using MACS2, bigWig tracks were generated. To examine the signals around the transcriptional start sites (TSS), deeptools was used to plot the heatmaps using the signals around 3 kb regions of the three datasets, with four sets: Housekeeping genes, Silenced genes, SpILT genes, and the testicular-expressed genes that are not detected in sperm. We analyzed previously published ATAC-seq libraries from mouse sperm (GSM2088376, GSM2088377, GSM2088378)^52^.

### Data availability

Next-generation sequencing data used in this study have been deposited at the NCBI Gene Expression Omnibus under the accession number GSE137490.

### Grouping transcripts based on expression dynamics

The abundance (tpm) of intact transcripts at each stage was calculated as a fraction of the maximum abundance reached during the developmental time course. A pseudo count of 0.001 was added before log transformation. CLARA method from the ‘cluster’ package was used in the clustering step, and 8 clusters gave the best separation. The data were clustered together, and then plotted by separating sperm and testis intact mRNAs and lncRNAs. Clustering results were visualized using Java Tree View 1.1.3.

### Gene Ontology analysis

Gene Ontology (GO) analysis was performed using clusterProfiler (version 3.12.0) package from Bioconductor (Release 3.9) {Yu et al., 2012, #43799}. Gene symbols were first converted to Entrez gene id. Enrichment analysis was performed by enrichGO function, using all genes detected in sperm and testis as background. Finally, we used dotplot function to visualize the GO results.

### miRNA target search and cumulative plots

miRNA target search was performed by miRanda^78^ using the -strict parameter. To compare the effects of miRNAs, cumulative plots were drawn in R using empirical distribution function, and p values were calculated by Kolmogorov–Smirnov test.

## Acknowledgments

We thank: D. Amador and the Interdisciplinary Center for Biotechnology Research facility and UR Genomics Research Center for help with the experiments, N. Chen and S. Pitnick for the cartoons of human and mouse sperm, and members of the Li and Au laboratories for discussions. This work was supported in part by National Institutes of Health grants K99/R00HD078482 to X.Z.L.

## Author Contributions

Y.H.S., A. W. and C.S. analyzed the data with input from K.F.A., and X.Z.L.; Y. H.S., and R.K.S. performed the experiments with input from K.F.A., and X.Z.L., and K.F.A., and X.Z.L. contributed to the design of the study, and all authors contributed to the preparation of the manuscript.

## Author Information

The authors declare no competing financial interests. Correspondence and requests for materials should be addressed to

## Supplemental Table Legends

**Table S1. Sperm purify quantification**

**Table S2. Sequencing Statistics**

(A) Mouse PacBio sequencing statistics.

(B) Human PacBio sequencing statistics.

(C) Small RNA sequencing statistics: reads and species.

(D) RNA-seq statistics: reads and species.

(E) PCR primers.

(F) Sperm intact long transcript (SpILT) information.

(G) Testicular sperm to cauda sperm RNAseq DESeq2 results, *q* < 0.1.

(H) MSUS (<u>M</u>aternal <u>S</u>eparation combined with <u>U</u>npredictable maternal <u>S</u>tress) sperm to control sperm RNAseq DESeq2 results, *q* < 0.1.

