## Supplemental File 1 for "Single-molecule long-read sequencing reveals a conserved selection mechanism determining intact long RNA and miRNA profiles in sperm"

**This PDF file includes:**

Caption for Additional Data Supplementary Table 1-2
Supplementary Fig. 1-4

**Other Supplemental Materials for this manuscript includes the following:**

Supplementary Table 1
Supplementary Table 2 (separate file)

Detailed information and statistics for the sequencing data used in this study.

**Table S1. Sperm purify quantification**

**Table S2. Sequencing Statistics (separate file)**

(A) Mouse PacBio sequencing statistics.

(B) Human PacBio sequencing statistics.

(C) Small RNA sequencing statistics: reads and species.

(D) RNA-seq statistics: reads and species.

(E) PCR primers.

(F) Sperm intact long transcript (SpILT) information.

(G) Testicular sperm to cauda sperm RNAseq DESeq2 results, $q < 0.1$.

(H) MSUS (Maternal Separation combined with Unpredictable maternal Stress) sperm to control sperm RNAseq DESeq2 results, $q < 0.1$.

### SUPPLEMENTARY FIGURE LEGENDS

**Supplementary Fig. 1. Defining intact transcripts in mouse sperm.** a) Bioanalyzer profiles of mouse testis, mouse sperm, and human sperm. b) Iso-seq library construction strategy. c) Potential artifacts that occur during Iso-seq library construction. d) Transcript detection of long read sequencing from mouse testis, mouse sperm, and human sperm. Rarefaction plot shows the number of transcripts detected (*y*-axis) as more reads were added to the analysis (*x*-axis). Different numbers of reads were randomly sampled from the successfully aligned reads, and the number of transcripts was calculated from the sampled reads. The black bar at each read count represents the range of detected transcripts from multiple sampling tests; the solid lines concatenate medians. The blue solid line represents the full-length matched transcripts only; red solid line represents transcripts detected with any match (full-length or partial). The fewer transcripts detected per increased number of reads sampled (i.e., the more the line flattens), the closer the analysis is to achieving complete detection (100% sensitivity). e) Novel transcript structures in sequencing from mouse sperm. Top to bottom: Genomic location; Ref-seq; PacBio reads; Illumina reads.

**Supplementary Fig. 2. Intact transcripts found in sperm.** a), Aggregated data for RNA-seq abundance on ribosomal protein-encoding mRNAs from sperm (*top*), and from testis (*bottom*) across 5´UTRs, ORFs, and 3´-UTRs. The x-axis shows the median length of these regions, and the y-axis represents the 10% trimmed mean of relative abundance. b) Testis RNAs are enriched for spermatogenesis functions. c) A scatterplot of SpILT abundance in control (2 biological replicates) vs MSUS conditions (4 biological replicates). Data analyzed were previously published[19]. Open circles represented SpILTs with significant changes ($q < 0.1$).

**Supplementary Fig. 3. Post-transcriptional selection for SpILTs.** a) Browser view of long reads from sperm at the *Crisp1* gene. No long reads at *Crisp1* gene were detected in testis. b) SpILT abundance in two biological replicates of testicular sperm (left) and cauda sperm (right). Tpm, transcripts per million. Pearson correlation coefficient (r) was computed. c) Box plot showing RNA abundance in purified round spermatids. Gold, SpILT. Purple, intact testicular RNAs that were not detected in sperm. In S3c and S3d, the analysis was made on previously published purified round spermatid RNA-seq data[50]. d) Correlation between SpILT abundance in mouse testicular sperm versus RNA abundance in purified round spermatids. Pearson correlation coefficient (r) was computed for mRNA and lncRNA separately. e) ATAC-seq footprint aggregate plots by distance to transcription start sites of SpILT genes, testicular-expressing genes whose transcripts are not detected in sperm, housekeeping genes and Germ-line silenced genes. Each row represents one transcript, normalized to sequencing depth.

**Supplementary Fig. 4. Post-transcriptional selection for sperm miRNAs.** a) Length distribution of total small RNAs and oxidized small RNAs, miRNAs are shown in gold. These piRNA reads with a peak of length at 30 nt are resistant to oxidation treatment. b) A scatterplot of anti-sense piRNA abundance mapping to each TE family in sperm vs. in round spermatids. Pearson correlation coefficient (r) was computed. c) A scatterplot of miRNA abundance from two biological replicates of testicular sperm. d) A scatterplot of miRNA abundance from two biological replicates of cauda sperm. e) miRNA binding sites are enriched within mRNAs that retained in human sperm. Plotted are cumulative distributions of SpILT abundance fold changes in sperm to those in round spermatids. We used top 10 most abundant miRNAs detected in human sperm small RNA-seq[76] and used MiRanda for the target scan[81]. f) Binding sites of miRNAs that increase in MSUS sperm are enriched within mRNAs that increase in MSUS sperm. Plotted are cumulative distributions of RNA fold changes in MSUS sperm to control sperm.

Table S1. Sperm purify quantification

| Table S1. Sperm purify, less than 1/10,000 contamination | |
|---|---|
| **Template** | **qPCR quantification of *Myh11*** |
| **Epididymis cDNA** | $1.26 \times 10^{-2}$ |
| **1/10 Epididymis cDNA** | $5.79 \times 10^{-3}$ |
| **1/100 Epididymis cDNA** | $6.80 \times 10^{-4}$ |
| **1/1,000 Epididymis cDNA** | $3.85 \times 10^{-5}$ |
| **1/10,000 Epididymis cDNA** | Not detected |
| **Purified sperm cDNA** | Not detected |

Sun et al, Fig. S1

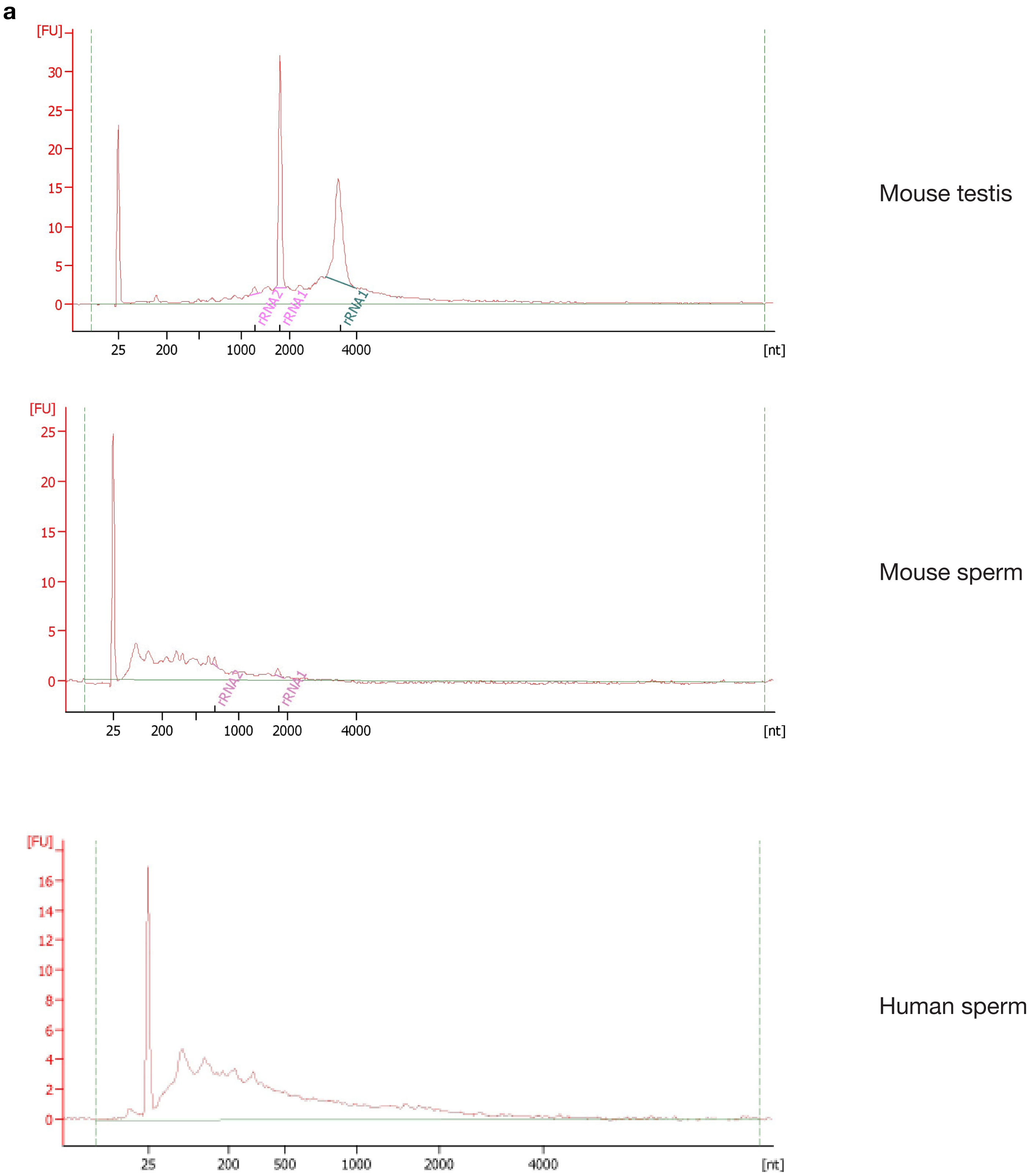

**b**

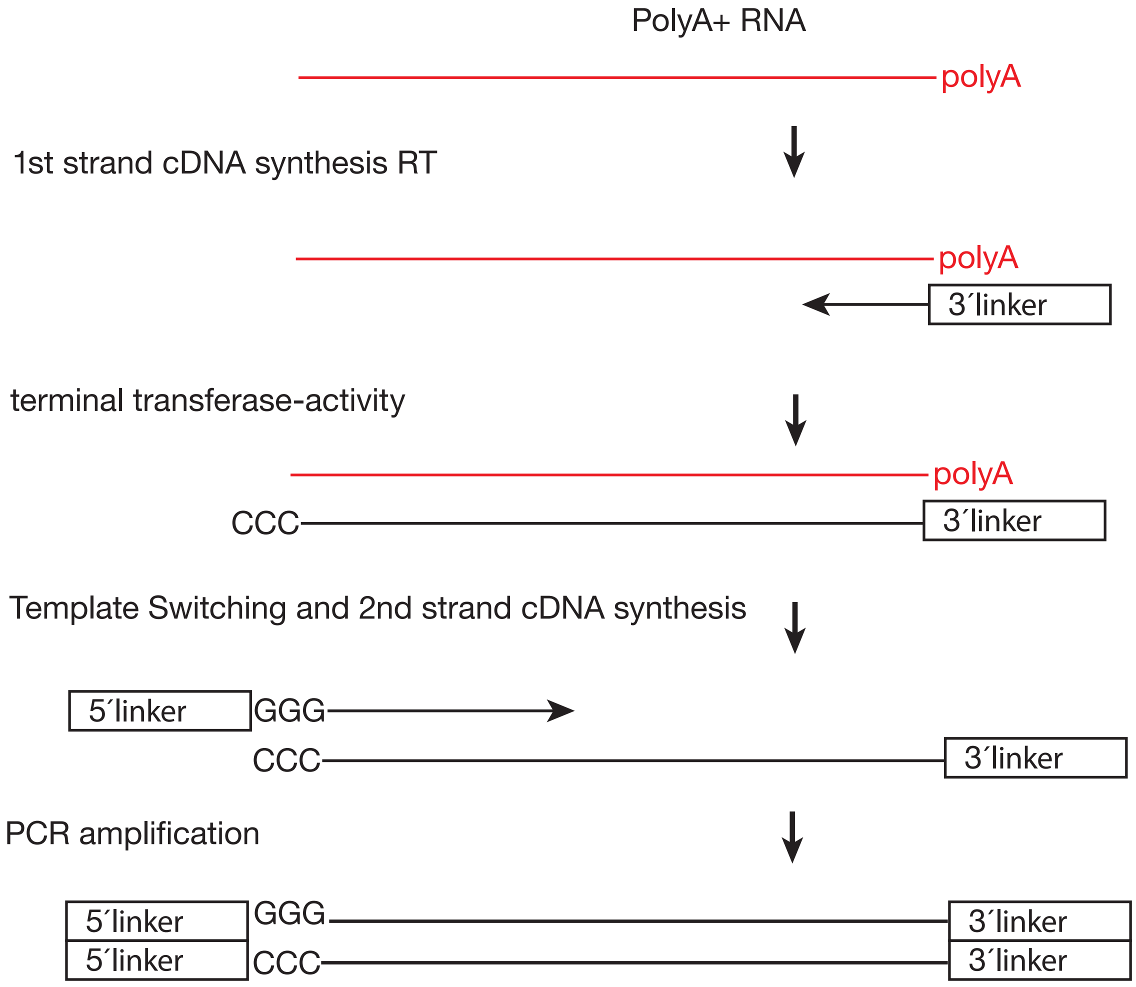
PolyA+ RNA
polyA
1st strand cDNA synthesis RT
polyA
3´linker
terminal transferase-activity
polyA
CCC
3´linker
Template Switching and 2nd strand cDNA synthesis
5´linker
GGG
CCC
3´linker
PCR amplification
5´linker
5´linker
GGG
CCC
3´linker
3´linker

**c**

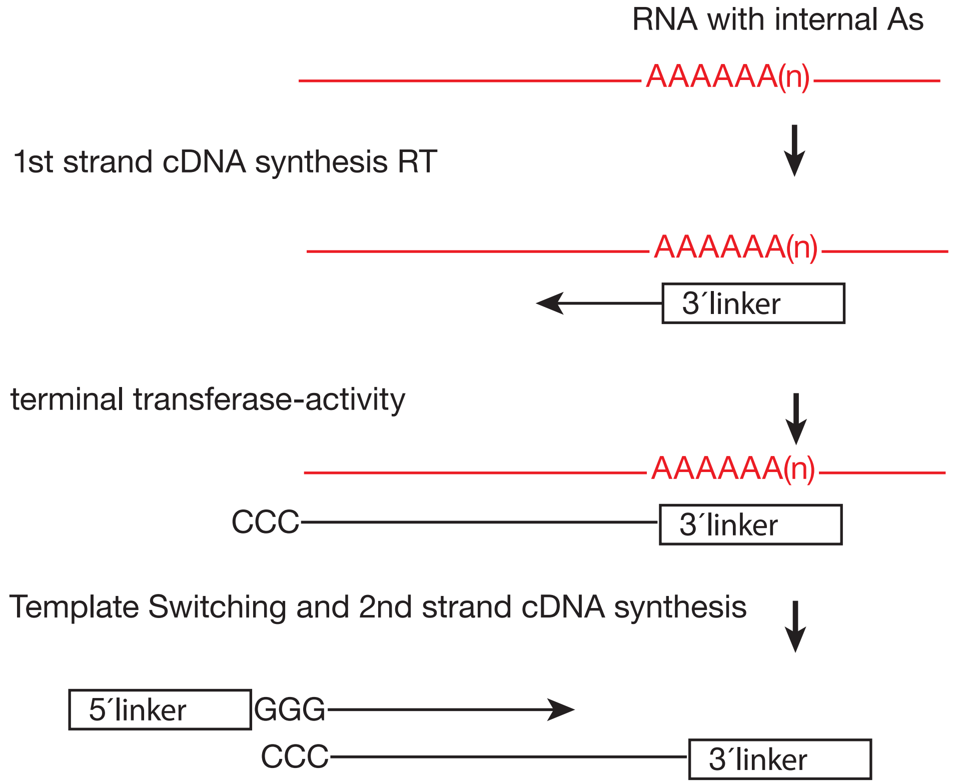
RNA with internal As
AAAAAA(n)
1st strand cDNA synthesis RT
AAAAAA(n)
3´linker
terminal transferase-activity
AAAAAA(n)
CCC
3´linker
Template Switching and 2nd strand cDNA synthesis
5´linker
GGG
CCC
3´linker

Sun et al, Fig. S1

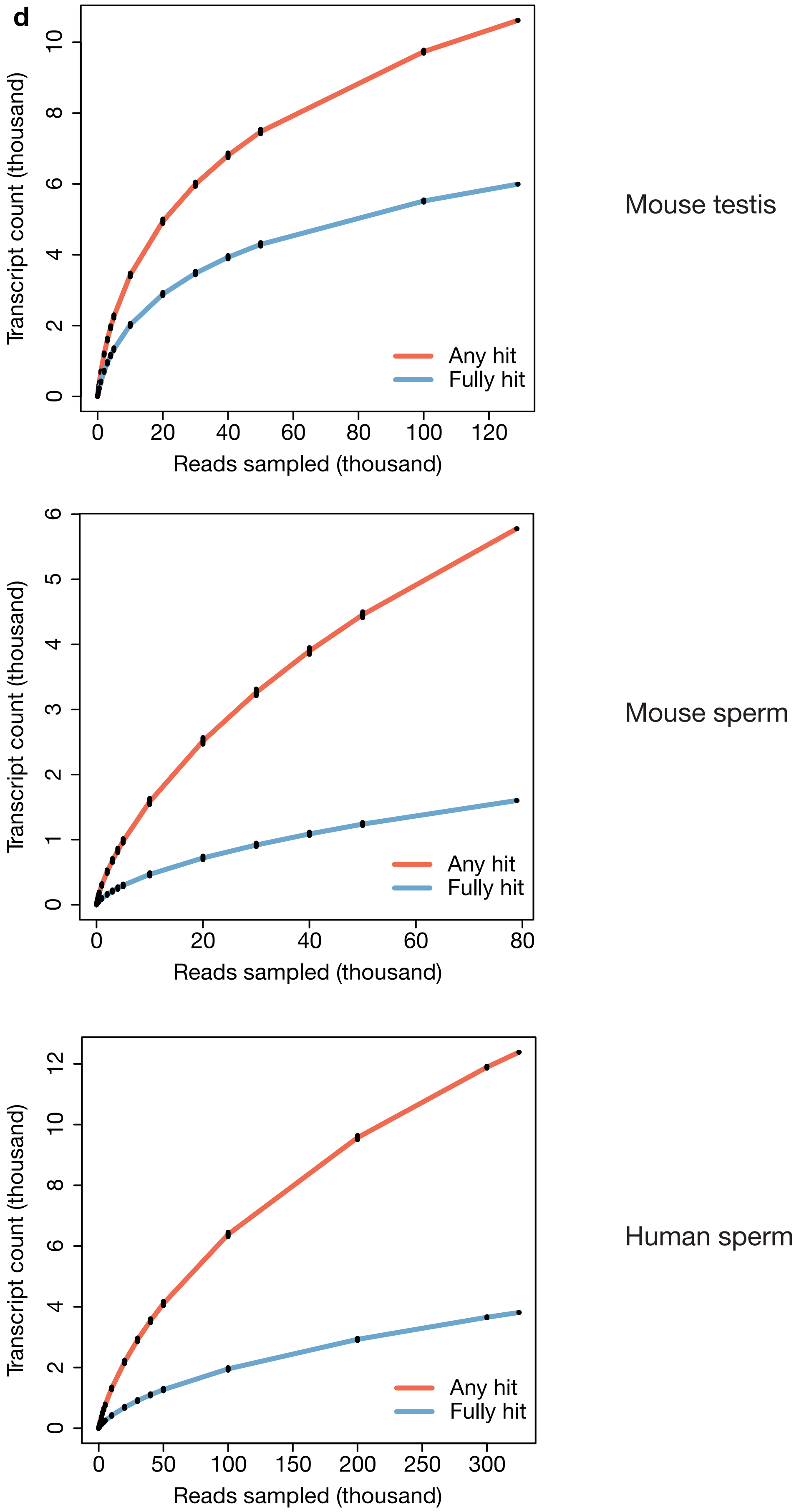

Sun et al, Fig. S1

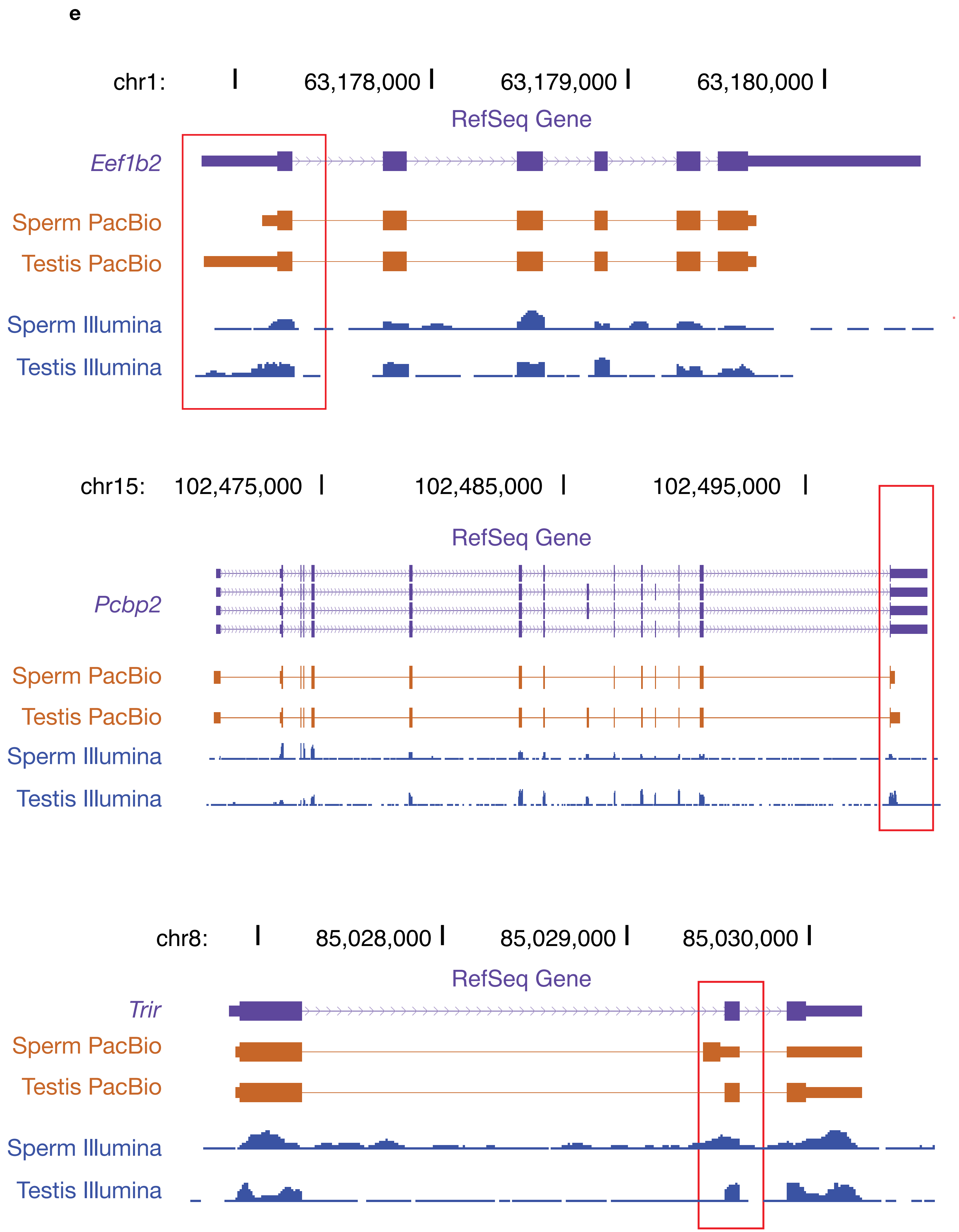

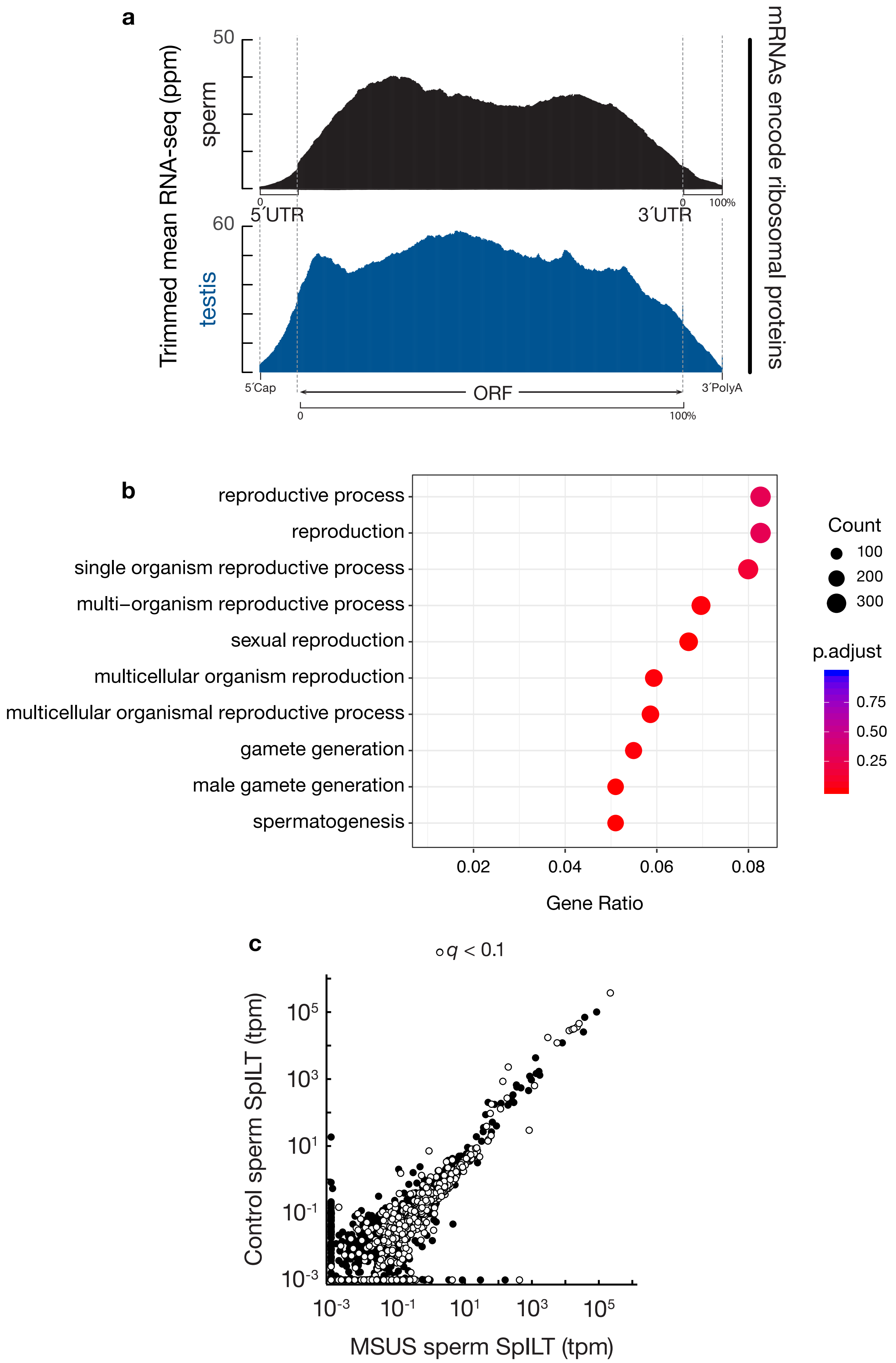
a
Trimmed mean RNA-seq (ppm)
50
sperm
60
testis
5´UTR
3´UTR
0
100%
5´Cap
ORF
3´PolyA
mRNAs encode ribosomal proteins
b
reproductive process
reproduction
single organism reproductive process
multi−organism reproductive process
sexual reproduction
multicellular organism reproduction
multicellular organismal reproductive process
gamete generation
male gamete generation
spermatogenesis
0.02
0.04
0.06
0.08
Gene Ratio
Count
100
200
300
p.adjust
0.75
0.50
0.25
c
q < 0.1
Control sperm SpILT (tpm)
MSUS sperm SpILT (tpm)
10^5
10^3
10^1
10^-1
10^-3

Sun et al, Fig. S3

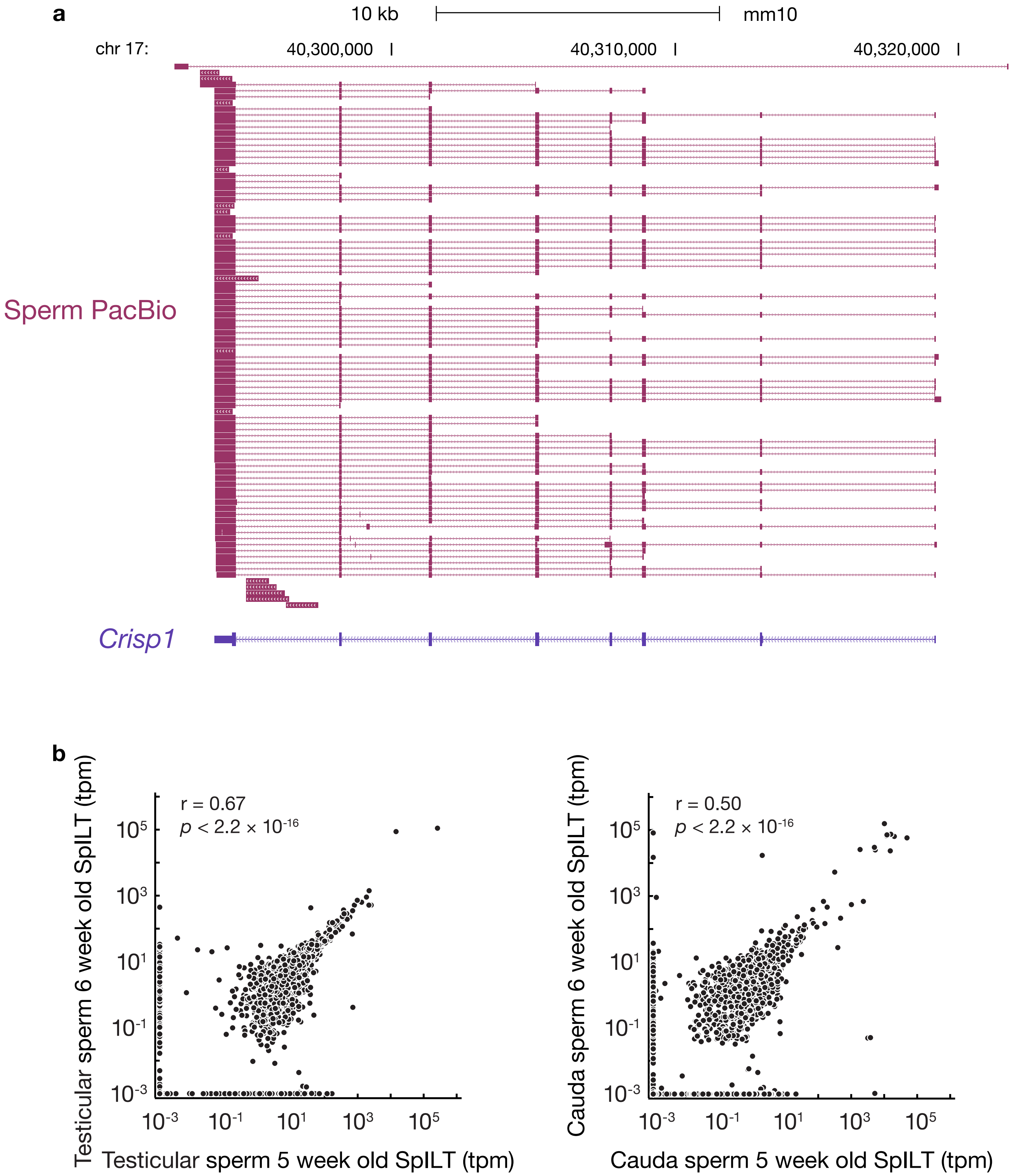

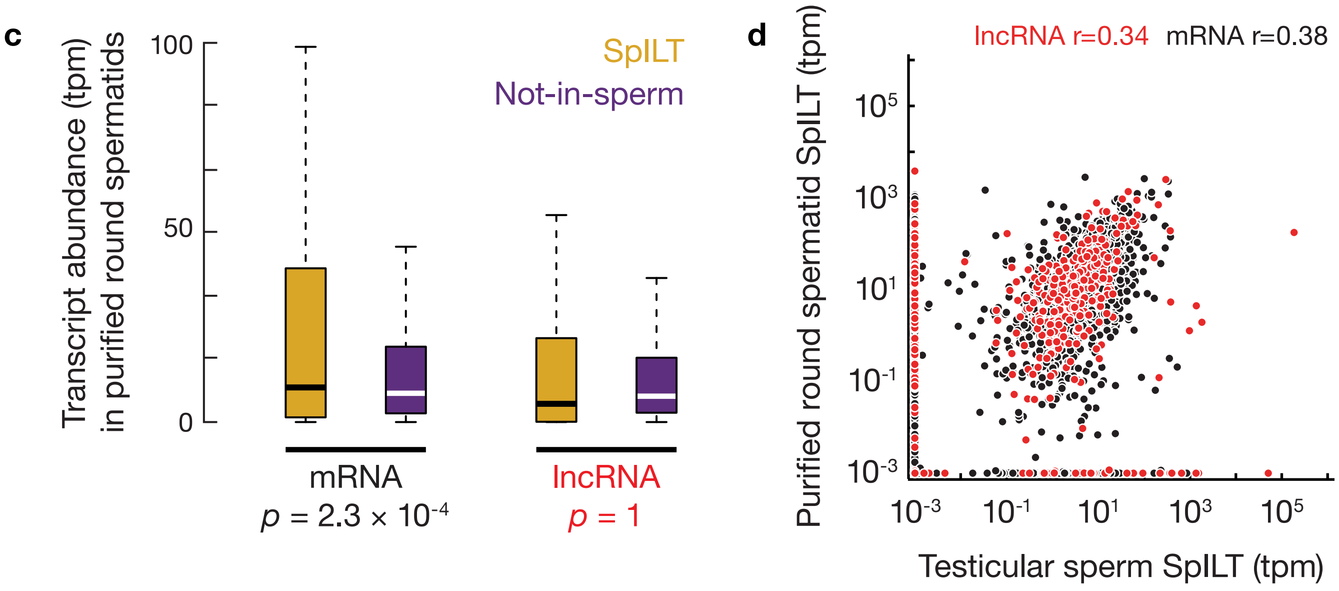
c
Transcript abundance (tpm) in purified round spermatids
100
50
0
SpILT
Not-in-sperm
mRNA
p = 2.3 × 10-4
lncRNA
p = 1
d
lncRNA r=0.34
mRNA r=0.38
Purified round spermatid SpILT (tpm)
10-3
10-1
101
103
105
Testicular sperm SpILT (tpm)

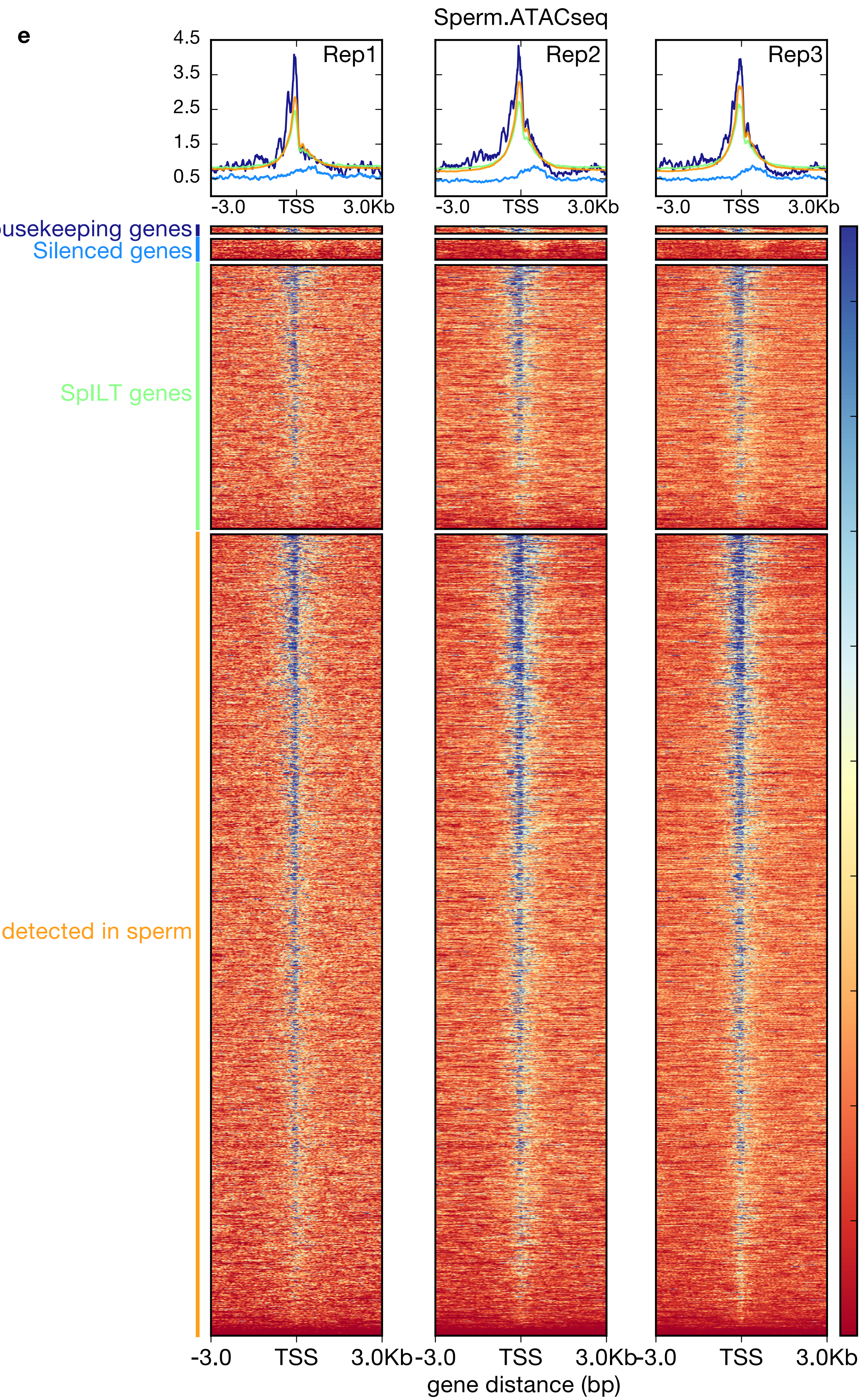
e
Sperm.ATACseq
Rep1
Rep2
Rep3
4.5
3.5
2.5
1.5
0.5
-3.0
TSS
3.0Kb
Housekeeping genes
Silenced genes
SpILT genes
Not detected in sperm
3.6
3.2
2.8
2.4
2.0
1.6
1.2
0.8
0.4
0.0
gene distance (bp)

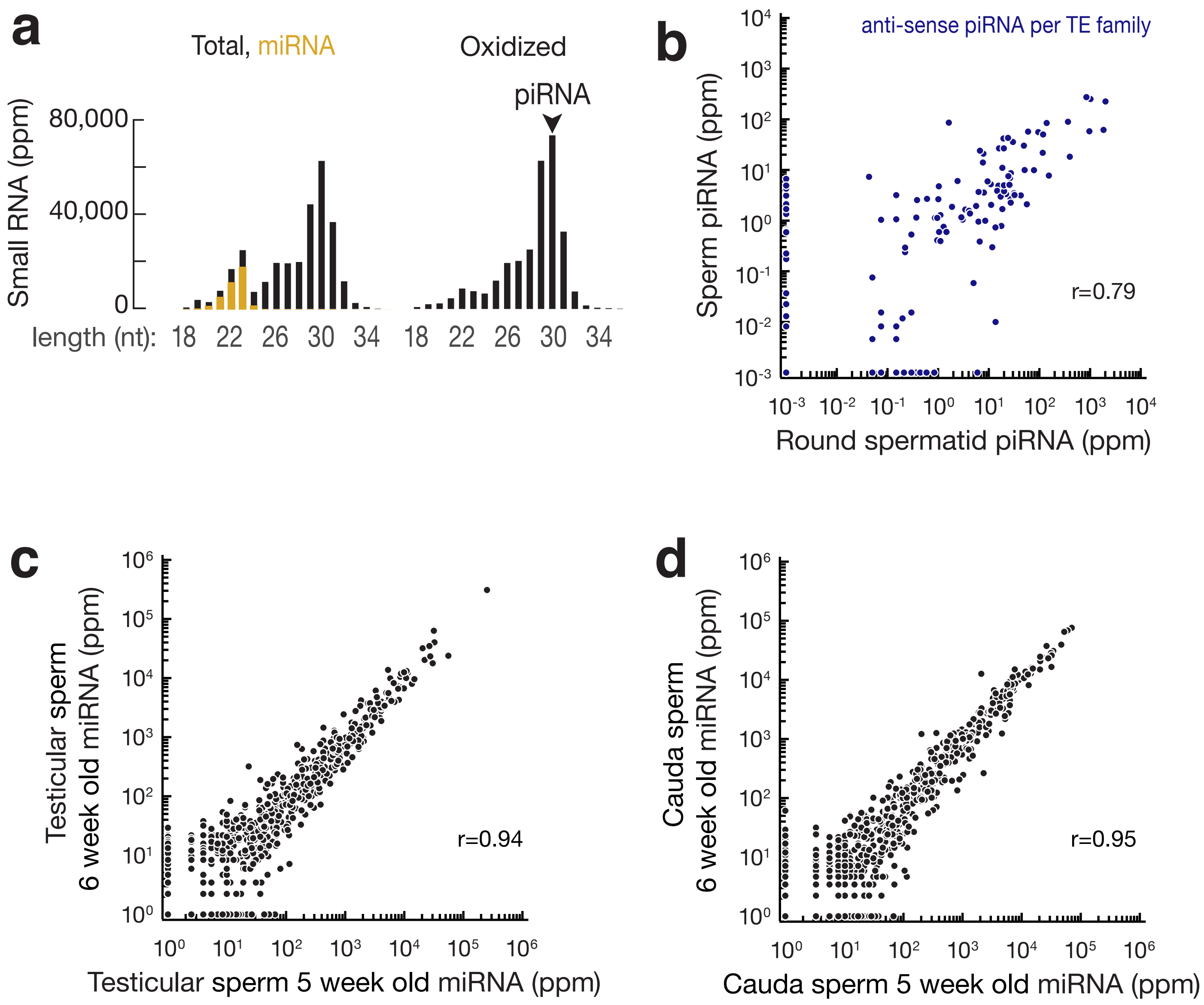
a
Total, miRNA
Oxidized
piRNA
Small RNA (ppm)
80,000
40,000
0
length (nt): 18 22 26 30 34 18 22 26 30 34
b
anti-sense piRNA per TE family
Sperm piRNA (ppm)
r=0.79
Round spermatid piRNA (ppm)
c
Testicular sperm
6 week old miRNA (ppm)
r=0.94
Testicular sperm 5 week old miRNA (ppm)
d
Cauda sperm
6 week old miRNA (ppm)
r=0.95
Cauda sperm 5 week old miRNA (ppm)

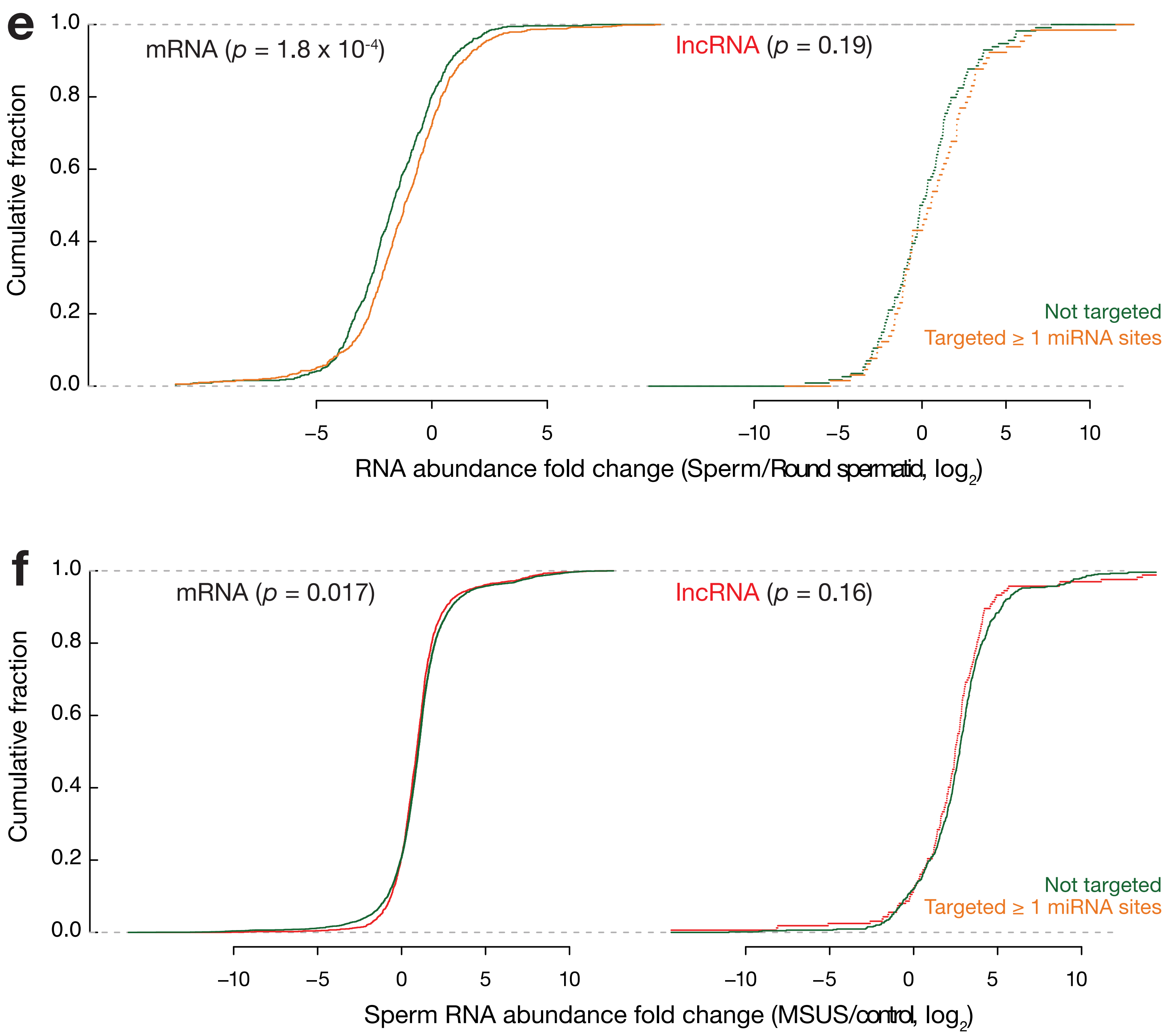
e
mRNA (p = 1.8 x 10-4)
lncRNA (p = 0.19)
Cumulative fraction
1.0
0.8
0.6
0.4
0.2
0.0
Not targeted
Targeted ≥ 1 miRNA sites
–5
0
5
–10
–5
0
5
10
RNA abundance fold change (Sperm/Round spermatid, log2)
f
mRNA (p = 0.017)
lncRNA (p = 0.16)
Cumulative fraction
1.0
0.8
0.6
0.4
0.2
0.0
Not targeted
Targeted ≥ 1 miRNA sites
–10
–5
0
5
10
–10
–5
0
5
10
Sperm RNA abundance fold change (MSUS/control, log2)
